# Identifying protein sites contributing to vaccine escape via statistical comparisons of short-term molecular dynamics simulations

**DOI:** 10.1101/2021.12.06.471374

**Authors:** Madhusudan Rajendran, Maureen C. Ferran, Gregory A. Babbitt

## Abstract

The identification of viral mutations that confer escape from antibodies is crucial for understanding the interplay between immunity and viral evolution. We describe a molecular dynamics (MD) based approach that scales well to a desktop computer with a high-end modern graphics processor and enables the user to identify protein sites that are prone to vaccine escape in a viral antigen. We first implement our MD pipeline to employ site-wise calculation of Kullback-Leibler divergence in atom fluctuation over replicate sets of short-term MD production runs thus enabling a statistical comparison of the rapid motion of influenza hemagglutinin (HA) in both the presence and absence of three well-known neutralizing antibodies. Using this simple comparative method applied to motions of viral proteins, we successfully identified *in silico* all previously empirically confirmed sites of escape in influenza HA, predetermined via selection experiments and neutralization assays. Upon the validation of our computational approach, we then surveyed potential hot spot residues in the receptor binding domain of the SARS-CoV-2 virus in the presence of COVOX-222 and S2H97 antibodies. We identified many single sites in the antigen-antibody interface that are similarly prone to potential antibody escape and that match many of the known sites of mutations arising in the SARS-CoV-2 variants of concern. In the omicron variant, we find only minimal adaptive evolutionary shifts in the functional binding profiles of both antibodies. In summary, we provide a fast and accurate computational method to monitor hot spots of functional evolution in antibody binding footprints.

## Introduction

In current attempts to prevent the spread of the Coronavirus disease 2019 (COVID-19) pandemic, caused by severe acute respiratory syndrome coronavirus 2 (SARS-CoV-2), clinicians and scientists have focused their efforts on the development of vaccines that are hoped to induce broad, long-lasting, neutralizing antibodies. However, the selective pressures imposed by the presence of the neutralizing antibodies in the host population can also drive the evolution of viruses towards adaptations that allow them to escape neutralization. For example, in the case of influenza viruses, immunity is provided by antibodies that target the hemagglutinin (HA), responsible for viral attachment and viral fusion to the host cell. However, these antibodies drive selection for amino acid substitutions in the HA, causing the influenza virus to rapidly evolve every year (1). Similarly, the antibody selection by the host immune system can also drive the emergence of new SARS-CoV-2 variants. Therefore, when developing vaccines that elicit antibodies against a broad range of strains, research efforts should also be aimed at identifying potential mutations that can facilitate viral escape from the neutralization effects of specific antibodies (2, 3).

The traditional approach to identifying these mutations is by passaging the virus in the presence of antibodies in a directed selection experiment, followed by validation of the variants that arise with neutralization assays. For example, in influenza viruses, escape mutant selection using a panel of monoclonal antibodies (mAbs) was used to identify the five major antigenic regions, Sa, Sb, Ca1, Ca2, and Cb (4-6). However, a significant drawback of this approach is that the directed selection typically only favors one of the many potential mutations that can escape a given antibody. Another approach is to test antibody binding to a panel of viral variants. In a typical 500 residue viral protein, there are about 10^4^ potential single amino acid mutants (7). Creating all individual mutants and then testing the mutants against the antibodies is an impossible task, thus causing researchers to limit themselves to exploring only a small portion of protein space (e.g., examining only mutations to alanine) (8, 9). Such studies cannot give a complete picture of the mutational spectra that can allow a virus to escape neutralization by a given antibody (10).

Functional evolutionary studies of viral vaccine escape are often supplemented with protein structural determination via x-ray crystallography or cryogenic electron microscopy. While structural biology can provide a static image of how an antibody physically contacts a viral protein, it cannot provide complete information regarding which amino acid sites are more prone to single replacement mutations (i.e. hot-spot sites or residues). Prediction of hot-spot residues is crucial as these sites on the antibody-antigen complex have a strong propensity to disrupt binding interactions within the antibody-antigen interface (11, 12). Recently, it has been demonstrated, via site-directed selection experiments and neutralization assays, that these hot-spot regions share a common biophysical feature. They all tend to harbor single amino acid sites that have significant large affects upon binding interactions in the antibody-antigen interface (13). Given this common feature of hot-spot residues, we hypothesize that some simple analyses of computer-based dynamics simulations of the antibody-antigen interface might help predict potential vaccine escape mutations before they happen, allowing for important functional context to real-time sequence-based surveillance of current and future pandemics.

Here we utilize a relatively simple method of comparative statistical analysis of molecular dynamic (MD) simulations developed by our lab (DROIDS - Detecting Relative Outlier Impacts in Dynamic Simulation or DROIDS 4.0) that employs a site-wise Kullback-Leibler (KL) divergence metric and a multiple test corrected two sample Kolmogorov-Smirnov (KS) test to successfully validate previously known sites of antibody escape in the influenza HA (13-17). We then utilized our site-wise comparative MD approach to identify potential sites prone to antibody escape in the spike protein of SARS-CoV-2. Specifically, in the omicron variant we were able to identify sites in the receptor binding domain (RBD) that support the binding efficiency to two general neutralizing antibodies and its competitive binding to the natural receptor, human angiotensin-converting enzyme 2 (hACE2). In summary, we present a method to accurately identify hot-spot residues that are prone to single point mutations with large functional effects upon the antibody-antigen binding interface and thus are likely pre-adapted to allow for vaccine escape. Identifying such residues *in silico* will be essential for pre-screening the antigenic consequences of viral genetic variations and designing better vaccines that induce long-term and broadly neutralizing antibodies against viral pathogens.

## Methods

### PDB structure, glycosylation, and model preparation

The protein structures used for the primary models for analyzing the molecular dynamics of antibody interactions with HA and the SARS-CoV-2-RBD are listed in Figure 1A. Any crystallographic reflections were removed along with any other small molecules used in crystallization. Hydrogens were added, and crystallographic waters were removed using pdb4amber (AmberTools18) (18). Glycosylation was deleted using the swapaa function in UCSF Chimera 1.14 (19). Predicted glycosylation were rebuilt for the Amber force field using the glycoprotein builder on the glycam.org webserver and the GLYCAM06j-1 force field (20, 21). Any missing loop structure in the files was inferred via homology modeling using the ‘refine loop’ command to Modeler within UCSF Chimera (22, 23). The globular head domain of the influenza HA is stable in its monomeric form. However, the stalk domain needs to be in a trimer to be stable (24). Therefore, MD simulations of PDB 4GMS was performed using trimmed (residues 57-270) monomeric form of the head domain. For MD simulations of the 3ZTJ and 4HLZ, the trimeric form of the stalk domain was used. The antibodies were trimmed, and only the heavy and light chains of the fragment antigen-binding (Fab) were used. Lastly, for the Omicron (B.1.1.529) variant simulations, we used the swapaa function to model the 15 RBD mutations onto 7OR9 (Figure 4B).

**Figure 1:**
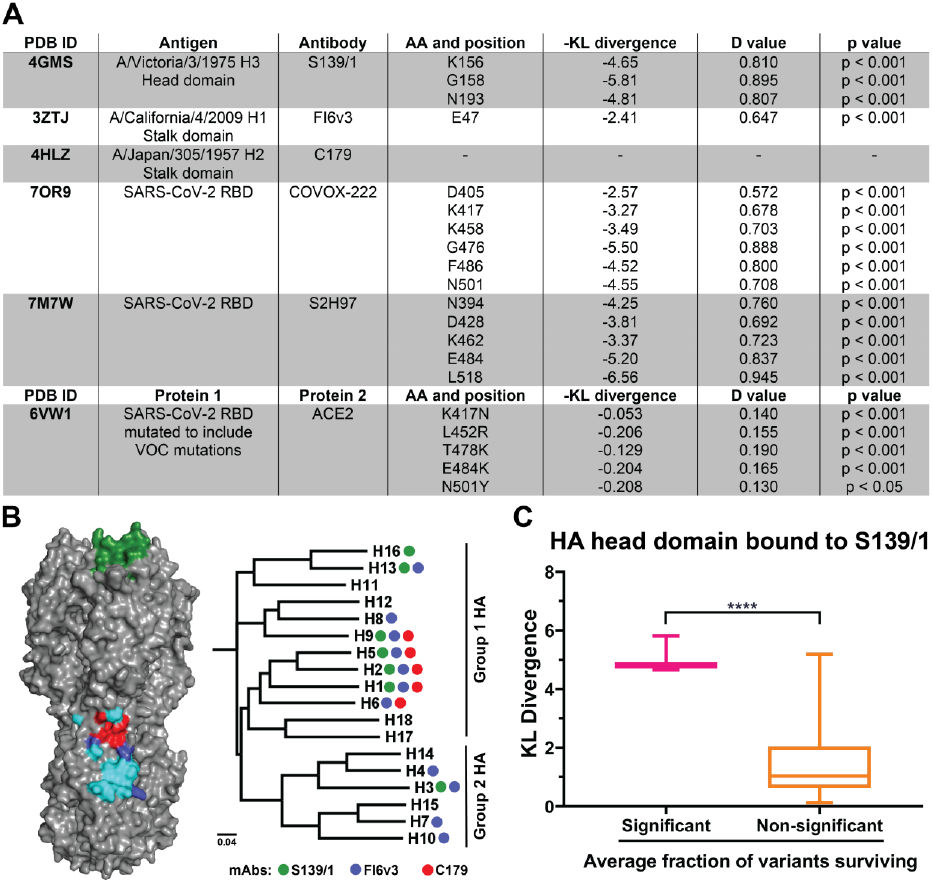
Description of PDB files and antibodies used in this study, and the comparison between KL divergence and average fraction of variants surviving. (A) Table summarizing the protein structure used for primary models for analyzing the molecular dynamics of antibody interactions with the influenza HA and SARS-CoV-2 RBD, and ACE2 interactions with wildtype and mutated SARS-CoV-2 RBD (includes VOC mutations). The table includes amino acid positions and the corresponding -KL divergence value denoting atomic fluctuations dampening for antibody-antigen/protein 1-protein 2 MD simulations. D value and the level of significance for the corresponding amino acid position are also given. (B) S139/1 (green), FI6v3 (blue), and C179 (red) epitopes are mapped onto the HA trimer, shown in grey (PDB 1RVX). Overlapping epitopes between FI6v3 and C179 are shown in cyan. Next to the HA trimer is a phylogenetic tree of HA subtypes. Circles denote reported antibody binding or neutralization against that subtype (C) Box and whisker plots showing the KL divergence values between amino acid sites that had significant average fraction of variants surviving and non-significant average fraction of variants in the presence of monoclonal antibody S139/1 (13). Only sites K156, G158, and N193 of the HA head domain had significant average fraction surviving viral particles.

### Molecular dynamics simulation protocols

All comparative molecular dynamics analysis via our DROIDS pipeline was based upon 100 replicate sets of 1 nanosecond accelerated MD runs (i.e., 100×1ns MD runs in each comparative state, e.g., unbound vs. bound). MD simulations were conducted using the particle mesh Ewald method via accelerated MD (pmemd.cuda) in Amber18 and the ff14SB protein and GLYCAM06j-1 force fields and implemented on two RTX 2080 Ti graphics processor units controlled via Linux Mint 19 operating system (21, 25-30). These replicate sets were preceded by energy minimization, 300 picoseconds of heating to 300K, and ten nanoseconds of equilibration, followed by random equilibration intervals for each replicate ranging from 0-0.5 nanoseconds. All protein systems were prepared via tLeAP (Ambertools18) and explicitly solvated, and charge neutralized with Na+ and Cl-ions in a Tip3P octahedral water box set to 12 nm beyond the surface of each protein with periodic boundaries (18, 31). All simulations were regulated using the Anderson thermostat at 300K and one atmospheric pressure (32). Root mean square atom fluctuations and atom correlations were conducted in CPPTRAJ using the atomicfluct and atomicorr commands (33).

### Comparative protein dynamics analyses with DROIDS 4.0 and statistical analyses

Comparative signatures of dampened atom fluctuation during antibody binding were presented as protein site-wise divergence in atom fluctuation in the antibody-bound versus unbound states for each viral target protein. Divergences were calculated using the signed symmetric Kullback-Leibler (KL) divergence calculation in DROIDS 4.0. Significance tests and p-values for these site-wise differences were calculated in DROIDS 4.0 using two-sample Kolmogorov-Smirnov tests with the Benjamini-Hochberg multiple test correction in DROIDS 4.0. The mathematical details of DROIDS 4.0 site-wise comparative protein dynamics analysis were published previously by our group and can be found here (14, 15). This code is available at our GitHub web landing: https://gbabbitt.github.io/DROIDS-4.0-comparative-protein-dynamics/, is also available at our GitHub repository https://github.com/gbabbitt/DROIDS-4.0-comparative-protein-dynamics.

## Results and Discussion

Using our comparative MD analysis pipeline (DROIDS 4.0), we successfully identified and validated previously known antibody escape sites in the head and stalk domains of the influenza HA. We then expanded the scope of our study to identify potential mutational sites prone to antibody escape in the spike protein of SARS-CoV-2 RBD and its recent omicron genetic variant.

Initially, we applied a Mann-Whitney U-test to compare KL divergence values, generated using MD simulations, to amino acid residues with significant average fraction of variants surviving and non-significant average fraction of variants surviving in the presence of S139/1, (Figure 1C) (13). In our comparative analyses of MD simulations, we found significantly higher KL divergence in the average fraction of variants surviving during directed selection (p < 0.001), suggesting that our KL divergence metric might prove as a useful quantitative measure in discriminating sites with mutations leading to vaccine escape. We then applied our approach to anti-HA antibodies with a wide range of breadth against different influenza strains. S139/1 is a one such monoclonal antibody known to neutralize both group 1(H1, H2, H5, H9, and H13) and group 2 (H3) influenza viruses (Figure 1B) (34). Crystallization studies have revealed that the antibody targets highly conserved residues in the RBD of the head domain (35) (Figure 1B). Comparative MD simulations of HA bound to S139/1 and unbound have revealed strong dampening of atom fluctuations occurring at sites K156, G158, and N193 of the HA, with -KL divergence values -4.65 (D = 0.810, p < 0.001), -5.81 (D = 0.895, p < 0.001), and -4.81 (D = 0.807, p < 0.001), respectively (Figure 1A and 2A). Directed selection using mAb S139/1 revealed that these three residues are sites of strong escape (Figure 2B) (13). As expected, the three amino acid residues with the highest negative KL divergence in our comparison of atom fluctuations from MD simulations also fall directly in the empirically determined physical binding footprint of the antibody. And they are the same three sites where previous works have selected escape mutants in the H1, H2, and H3 HAs (34, 35). Residue 156 is a part of the 150 loop of the influenza HAand forms electrostatic interactions by inserting itself into the acidic pocket in the Fab formed by residues Glu^H35^ and Glu^H50^. Another 150 loop binding determinant is residue 158, which has the largest dampening of atomic fluctuations. Gly158 is closely stacked on the light chain complementarity determining region 3 (LCDR3) of S139/1 and is further stabilized by a main chain hydrogen bond between S159 and Asn^L92^. Any mutations at these positions will cause clashes at the antibody interface. S139/1 also recognizes members of the 190 helix. In the residues of 190 helix, Asn193 plays a significant role in antibody recognition and is buried by heavy chain complementarity determining regions (HCDR) 1-3 of S139/1 (34). Furthermore, it should be noted that site 193 was known to interact with the host receptor molecule (sialic acid moiety), suggesting the contribution of this residue to receptor binding of HA (36, 37). Therefore, isolates with mutated residues at site 193 have reduced viral fitness due to their inability to bind to the host receptors (38). Selective pressure created by S139/1 on residue 193 has other secondary effects on the virus, such as the reduced binding activity of the influenza virus.

**Figure 2:**
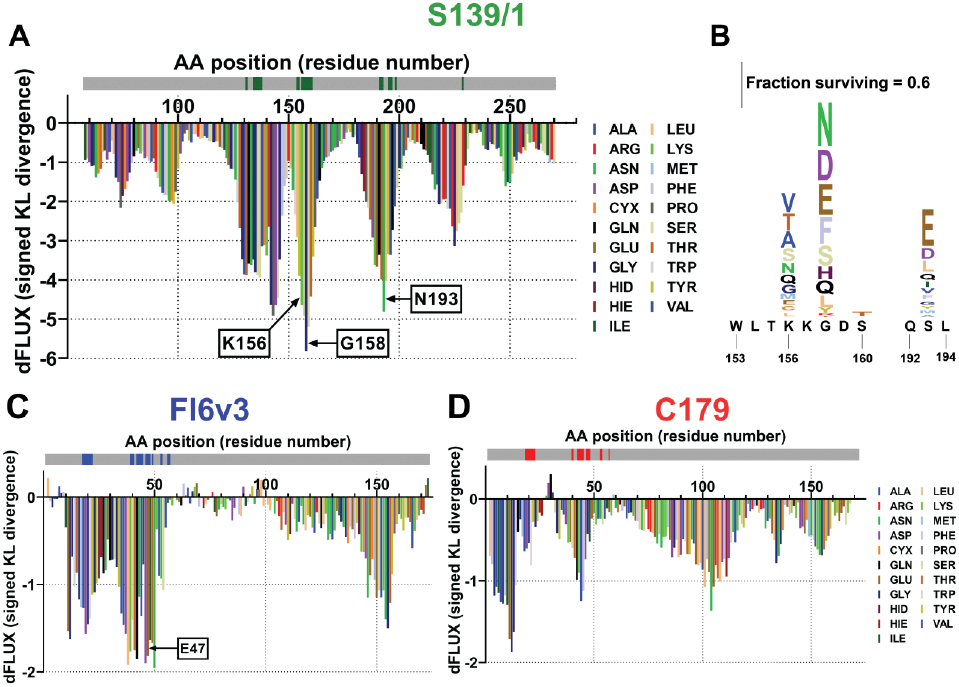
Epitope prediction and validation of hot-spot residues of the influenza hemagglutinin (HA) in the presence of neutralizing antibodies. Sequence positional plotting of dampening of atom motion on the influenza HA head domain by (A) S139/1 and on the HA stalk domain by (C) FI6v3 and (D) C179. The sequence profile of the -KL divergence between S139/1 and the head domain produces strong negative peaks in (A) at K156, G158, and N193. A modest negative peak is observed in the stalk domain in (C) at E47 in the presence of FI6v3. HA1 numbering is used to identify the amino acid positions in (A). HA2 numbering is used to identify the amino acid position in (C) and (D). The grey bar on top of the -KL divergence plots denotes the HA amino acid backbone with the location of (A) S139/1 epitopes shown in green, (C) FI6v3 epitopes in blue, and (D) C179 epitopes in red. (B) Logo plots of S139/1 show the amino acid position that has the largest effect. Letter heights are proportional to the excess fraction of virions with that mutation that survive the antibody, as indicated by scale bars. The logo plot was prepared using deep mutational scanning data from Doud, M.B. *et al*. (13).

Lastly, with mAb S139/1, atomic fluctuation dampening is also seen in residues 133-134 of the influenza HA (Figure 2A). These residues are part of the 130 loop of the HA, and make a significant portion of the RBD of the influenza virus. Lastly, in addition to atomic fluctuation dampening at residues 133-134 and at residues 156/158, we also see dampening at 145-147 (Figure 2A). Upon further examination, there lie no epitopes of S139/1 between residues 145-147. Therefore, we believe that this dampening is an artifact caused purely by the residue’s location between two epitope sites (130 loop and 150 loop) of S139/1. In summary, the three sites determined to have the largest physical effect upon antibody binding in our simulations (i.e. showing the largest negative KL divergence) are exactly the same three sites most likely to evolve antibody escape under directed selection experiments.

We also chose two broad antibodies that target the stalk domain of the HA (FI6v3 and C179) to further validate our computational approach to identify potential sites of escape. FI6v3 was first isolated by high-throughput screening of immortalized antibody-secreting cells and was found to bind to both group 1 (H1, H2, H5, H6, H8, H9, and H13) and group 2 (H3, H4, H7, and H10) viruses (Figure 1B) (39). Antibody C179 was first isolated from a mouse that had been immunized with the H2N2 virus and was later found to cross-neutralize H1, H2, H5, H6, and H9 subtypes (Figure 1B) (40). Both FI6v3 and C179 have epitopes that lie in the stalk domain and are known to interfere with membrane fusion (Figure 1B). Our MD simulations do not reveal dampening of atomic fluctuations in the stalk domain in the presence of FI6v3 or C179. When we look at the difference in atomic fluctuations of the stalk antibodies on the same scale as S139/1, we only see a few sites with strong dampening (Figures 2C and 2D).

Antibody selection revealed very similar results as our MD simulation, with only few sites in the stalk domain with slightly increased fraction of variants surviving mAbs C179 and (13). As a result, the authors concluded that the stalk domain is less capable of escaping antibodies by single mutations (13). In the presence of FI6v3, both directed selection and MD simulation show a small bump at site 47 (KL = - 2.41, D = 0.647, p < 0.001) that is of importance (Figures 1A and 2C) (13). Mutational studies have demonstrated that the introduction of E47R in the stalk domain has increased the resistance to FI6v3 (41). Like other studies, we also conclude that the HA stalk is intolerant of mutations, and confirmed by us here due to the lack of sites with dampened fluctuations. The absence of sites with large divergences in atomic fluctuations in our comparative MD simulations would seem to confirm that the anti-stalk antibodies studied here readily target sites with high mutational tolerance as suggested by other site directed experiments (42). Another possible explanation could be that the binding energetics at protein-protein interfaces can be asymmetrically distributed across all sites, thus preventing us from identifying the mutational tolerant sites (11, 12).

We implemented our computational approach to identify potential hot spot residues in the RBD of the spike protein of SARS-CoV-2 in the presence of two recently published antibodies against the virus: COVOX-222 and S2H97 (Figure 3A). COVOX-222 is known to bind to different residues than S2H97 and is known to neutralize strains P.1 (Gamma) from Brazil, B.1.351 (Beta) from South Africa, and B.1.1.7 (Alpha) from the United Kingdom (43). Termed “super-antibody,” S2H97 is known to bind with high affinity across all sarbecovirus clades and prophylactically protects hamsters from viral challenge (44, 45). In the presence of COVOX-222, we see the most dampening of atomic fluctuations at residues G476 (KL = -5.50, D = 0.888, p < 0.001), F486 (KL = -4.52, D = 0.800, p < 0.001), and N501 (KL = -4.55, D = 0.708, p < 0.001) of the RBD (Figure 1A and 3B). We see modest amount of dampening at residues D405 (KL = - 2.57, D = 0.572, p < 0.001), K417 (KL = -3.27, D = 0.678, p < 0.001), and K458 (KL = -3.49, D = 0.703, p < 0.001) of the RBD (Figure 1A and 3B).

**Figure 3:**
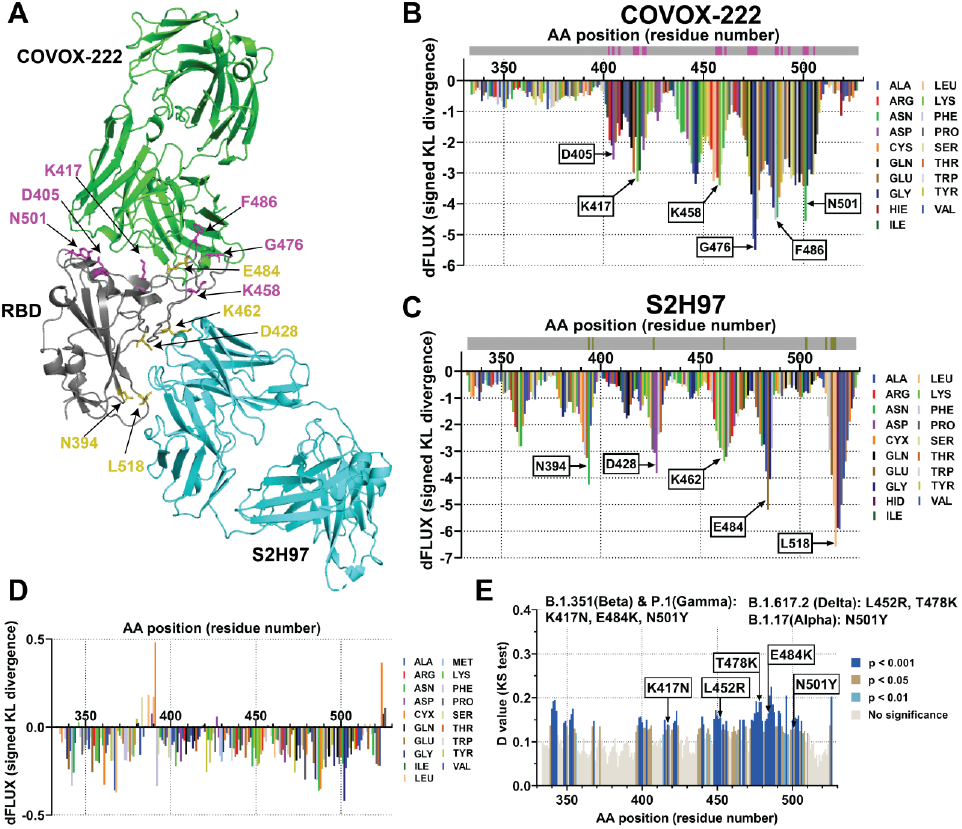
Identification of hot-spot residues that escape neutralizing antibodies and the importance of the variants of concern mutations. (A) Crystal structure of COVOX-222 (green, PDB 7OR9) and S2H97(cyan, PDB 7M7W) superimposed onto the structure of RBD of SARS-CoV-2 (grey, PDB 7M7W) (43, 44). COVOX-222 epitopes with the greatest KL divergence dampening are highlighted in pink, and S2H97 epitopes with the greatest KL divergence dampening are highlighted in olive. Sequence positional plotting of dampening of atom motion on the RBD of SARS-CoV-2 by (B) COVOX-222 and (C) S2H97. Amino acid positions with moderate to modest dampening of atomic fluctuation are identified in (B) and (C). The grey bar on the top of the KL divergence plots denotes the RBD domain amino acid backbone with the location of COVOX-222 epitopes shown in pink and the S2H97 epitopes shown in olive. (D) The site-wise KL divergence profiles showing the dampening of atom motion between the wildtype and mutated RBD of the spike protein of SARS-CoV-2 in the presence of human angiotensin-converting enzyme 2 (ACE2). The wildtype RBD was computationally mutated to include mutations from the variants of concern (VOC). (E) Multiple test-corrected two sample KS tests of significance for the impact of the mutations are also shown. Amino acid positions that correspond to the VOC mutations are highlighted and show higher level significance than other positions. Mutations that correspond to the four VOC in the RBD of spike protein are noted on top of (E).

**Figure 4:**
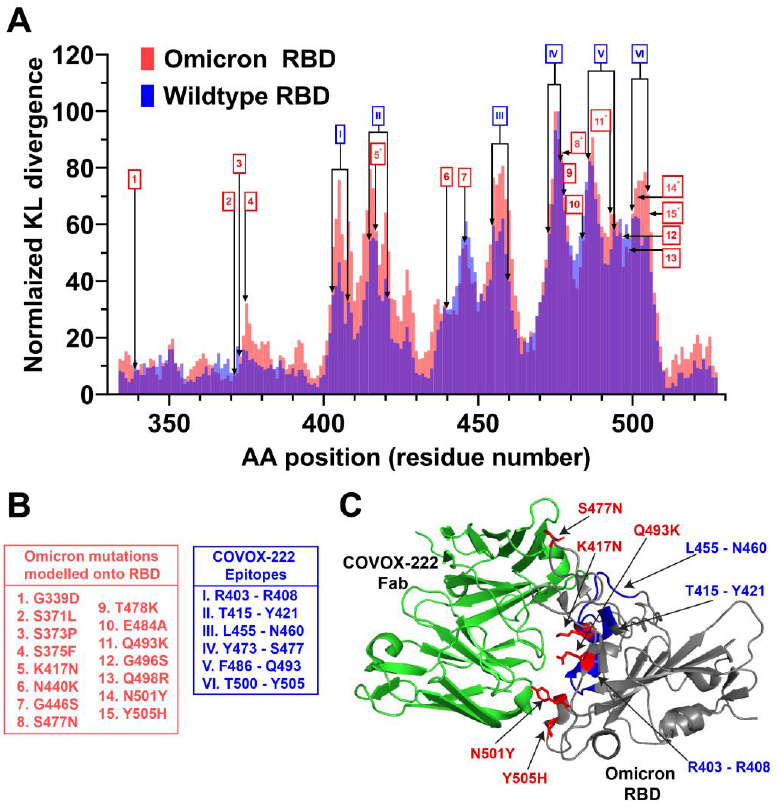
MD simulations with the Omicron variant reveal sites that promote similar binding affinity to hACE2. (A) Sequence positional plotting of the normalized dampening of atom motions on the Omicron RBD (red) and the wildtype RBD (blue) by monoclonal antibody COVOX-222. KL divergence values for normalization of the wild type RBD were obtained from Figure 3B, and for the Omicron RBD were obtained from supplemental figure 1A. The omicron RBD mutations are labeled in red (1-15). The sites corresponding to the COVOX-222 epitope on the wildtype RBD on labeled in blue (I – VII). The amino acid residues that correspond to the numbers and roman numerals are listed in (B). In the omicron RBD, we see an increase in atomic fluctuation peaks at several of the epitope sites. Sites that overlap between the omicron mutations and COVOX-222 epitope (shown by *) are sites of ACE2 interaction. (B) List of omicron mutations modeled onto RBD, and the list of COVOX-222 epitopes. (C) Crystal structure of COVOX-222 Fab (green) superimposed onto the structure of the Omicron RBD (grey) (PDB 7NXA). Some of the amino acid residues that correspond to the COVOX-222 epitopes, which show an increase in atomic fluctuation dampening, are shown in blue. Several of the amino acid residue that correspond to omicron mutations with increased atomic fluctuation dampening, which are sites of ACE2 interactions, are shown in red.

It should be noted that the residues picked up by our computational method are part of the epitopes of COVOX-222. Of all the residues identified by the comparison of MD simulation, the residues with the greatest dampening of atomic fluctuation have a higher chance of being classified as hot-spot residues. They are more prone to mutate, thus allowing the virus to escape COVOX-222. Residue 417, one of the residues with moderate dampening of atomic fluctuation, makes a weak salt-bridge interaction with the HCDR3 residue E99 of COVOX-222 (43). And residue N501, one with residues with the most dampening of atomic fluctuations, is known to interact with LCDR1 residue P30 of the antibody via a stacking interactions (43, 46). In the case of the mAb S2H97, we see a moderate dampening of atomic fluctuations at sites D428 (KL = -3.81, D = 0.692, p < 0.001) and K462 (KL = -3.37, D = 0.723, p < 0.001). Sites N394 (KL = -4.25, D = 0.760, p < 0.001), E484 (KL = -5.20, D = 0.837, p < 0.001), L518 (KL = - 6.56, D = 0.945, p < 0.001) have the most dampening of atomic fluctuations (Figure 1A and 3C). Four of the five sites, except E484, fall in the epitope footprint of S2H97 (44). Like mAb COVOX-222, we predict that the sites with the most dampening of atomic fluctuations (i.e. negative KL divergence) are more prone to functionally evolve under the selection pressure of the vaccine, thus allowing the virus to potentially escape the binding of S2H97. Even though, E484 does not fall in the S2H97 footprint, it has been shown that the mutation of this site is known to enhance immune escape form neutralizing antibodies, and also increase affinity to hACE2 (47).

Many of the residues identified as potential sites of escape, either in the presence of COVOX-222 or S2H97, overlap with mutations seen in the variants of concern (VOC). At present, there are mainly five kinds of VOC: Alpha, Beta, Gamma, Delta, and Omicron. We employed comparative MD simulation of the SARS-CoV-2 RBD bound to hACE2 (PDB 6VW1 trimmed to only include the viral RBD) modeled with and without the recently arising VOC mutations in the alpha to delta variant strains. Three of the VOC mutations (417, 484, and 501) were identified as potential escape sites using our approach (Figure 1A, 3B, and 3C). In addition to the three sites, we confirmed (i.e., 417, 484, and 501), we also saw significant differences in the binding atomic fluctuations at the positions 452 (KL = -0.206, D = 0.155, p < 0.001) and 478 (KL = -0.129, D = 0.190, p < 0.001) of RBD (Figures 1A, 3D and 3E). Investigation with pseudoviruses possessing RBD mutations harbored by VOC demonstrated that plasma neutralizing activity of vaccinated individuals showed a one-fold to three-fold significant decreases against E484K, N501Y, or the K417N + E484K + N501Y triple mutant (48). These results confirm what we have observed using our comparative MD analysis (Figures 1A and 3E). Furthermore, evidence from clinical trials on the impact of VOC on vaccine efficacy confirms what we have observed as well. For example, the ChAdOx1 nCoV-19 vaccine and the single-dose JNJ-78436735 (Johnson & Johnson/Janssen) vaccine have reduced vaccine efficacy by 10.4% and 57%, respectively, against the B.1.351 variant, which contains K417N, E484K, and N501Y mutations (49-51). Therefore, when evaluating vaccine efficacy and the effect of neutralizing antibodies against the SARS-CoV-2 virus, the focus should be given to amino acid positions that are prone to mutate and escape the antibody, as these positions might cause the emergence of new VOCs.

Lastly, we also modeled the most recent VOC, Omicron. The omicron RBD is known to harbor 15 different mutations, with several mutations linked to greater transmissibility, lower vaccine efficiency, and increased risk of reinfection (Figure 4B) (52). To further understand the evolution of the variant, we ran comparative MD simulations with the Omicron variant bound and unbound to mAb COVOX-222. In the wildtype RBD we see a dampening of atomic fluctuation at residues that correspond to the antibody footprint (Figure 4A, 4B). Interestingly, in the omicron RBD, we see some increase in the atomic fluctuation dampening at several of the residues that correspond to the COVOX-222 epitopes (Figure 4). Furthermore, some of the mutations that correspond to the omicron RBD overlap with the COVOX-222 antibody footprint. These sites include K417N (KL = -2.60, D = 0.631, p < 0.001), S477N (KL = -3.83, D = 0.762, p < 0.001), Q493K (KL = -2.84, D = 0.762, p < 0.001), N501Y (KL = -3.25, D = 0.737, p < 0.001), and Y505H (KL = -3.16, D = 0.77, p < 0.001) (Figure 4, Supplemental figure 1). Upon closer examination, the residues that overlap between the antibody footprint and the omicron mutations, are sites where the hACE2 is known to interact with the RBD. As a result, the increase in atomic fluctuation dampening might not be because of the direct evolution of stronger binding to the antibody, but also due to the evolution of increased binding affinity to hACE2. Other studies have shown that the mutations in the omicron variant increase the number of salt bridges and hydrophobic interactions between RBD and hACE2, resulting in a higher binding efficiency to hACE2 (53). Additionally, it has also been shown that the structural changes in the RBD domain, caused by the mutations, reduce the antibody interactions (54, 55). We present a similar analysis of the functional binding of the omicron variant with S2H97 (Supplemental Figure 2) which also indicates minimal overall effects of the genetic changes on potential antibody neutralization, with the exception of E484A, T478K and S477N sites, in which omicron appears to have lost interaction with S2H97. In summary, we observe that introduction of mutations sites in the omicron RBD have only slightly altered the COVOX-222 and S2H97 interactions with the viral RBD. Most importantly, our computational method appears to be an effective way to quantify these changes.

Directed selection experiments followed by confirmation through neutralization assays are the classical approach for identifying the location of the hot-spot residues. Not only are the selection experiments time consuming, they only identify one of many mutations that escape an antibody (56). Compared to the classical method, our simple computational approach can identify the locations of sites prone to vaccine escape in a matter of days. Additionally it identifies positions of all amino acids that could mutate to escape the antibody in a single *in silico* experiment. And compared to other computational tools, our MD-based approach does not require large and potentially biased machine learning training data sets. Furthermore, the *de novo* prediction is based on a given experimental structure, which enables an unprecedented close synergy between our computational approach and existing laboratory methods for identifying potential routes of viral evolution leading to enhanced transmissibility and vaccine escape.

Additionally, some of the other tools used in epitope prediction involve sequence-based methods, machine-learning methods, and structure-based methods. The epitope surface accessibility to antibody binding is generally used by the sequence-based methods (57). The availability of antigen sequence is crucial for the sequence-based method; however, the predicted epitope residues are not grouped into the corresponding epitopes (58). The machine learning (ML) based epitope prediction methods include several steps: a collection of datasets with clean and comprehensive data, extraction of antigen features of the sequence (e.g., physicochemical properties, evolutionary information, amino acid composition), and training the model using ML algorithms (59). Some of the commonly used ML tools for epitope prediction are ABCPred (uses artificial neural network), COBEpro (uses support vector machine), EPSVR (uses support vector regression method), and BepiPred (based on random forest algorithm) (57, 60-62). And lastly, structure-based methods identify epitopes by antigen structure and epitope-related propensity scales, including specific physicochemical properties and geometric attributes (63, 64). As the evolution of antibody-antigen interactions seem largely driven by single-site mutations with large functional effects and because antibody-antigen binding has quite obvious biophysical impacts on molecular dynamics, as would the formation of any protein complex, we offer a simple and effective alternative computational method to studying vaccine escape *in silico* using readily available MD simulation software combined with a classic and easily interpretable approach to the statistical comparison of distributions of local atom fluctuations on virus proteins in the presence/absence of antibodies.

In summary, we have computationally identified hot-spot residues known to mutate in a viral antigen in the presence of a neutralizing antibody. We first validated our approach using the influenza spike protein and three well-characterized antibodies against the influenza HA (13). We then implemented our approach to identify sites in the RBD of SARS-CoV-2 that are known to mutate in the presence of neutralizing mAbs S2H97 and COVOX-222 (43, 44). We further identified that residues known to mutate in the presence of the antibody overlap with the mutations seen in the VOC. And lastly, we also identified sites in the omicron variant mutations that enhance binding efficiency to hACE2. While the method described here is not a complete substitute for laboratory-based methods we believe it can greatly complement these methods by allowing time and cost-saving through the computational prescreening of the underlying biophysics that may drive outcomes of many potential lab experiments. Determining which viral mutations escape from antibodies will be crucial for designing future therapeutics and vaccines and assessing future antigenic implications of ongoing viral evolution.

## Author contributions

M.R wrote the manuscript, designed, performed the experiments, and analyzed the data. G.A.B developed the code, revised and edited the manuscript, designed the experiments, and analyzed the data. M.C.F edited the manuscript and analyzed the data.

## Acknowledgements

We acknowledge Dr. Ernest Fokoue (School of Mathematical Sciences, Rochester Institute of Technology) for his consultation on the prior development of our method. We also acknowledge Dr. Feng Cui and Dr. André O. Hudson (Thomas H. Gosnell School of Life Sciences, Rochester Institute of Technology) for helpful comments regarding this work.

---

**Supplemental Figure 1:**
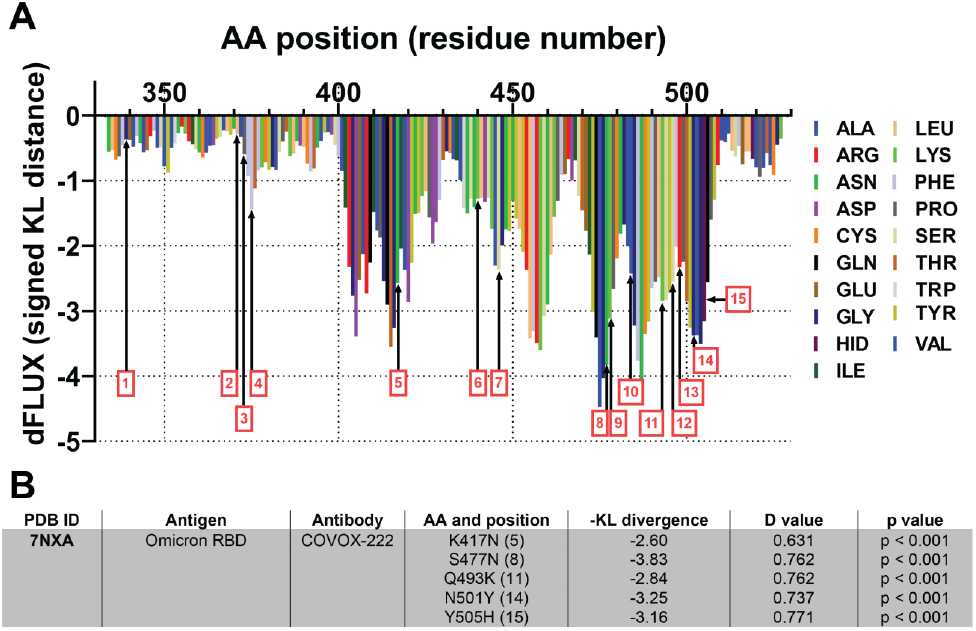
Molecular dynamic simulation of the omicron variant. (A) Sequence positional plotting of dampening of atom motion on omicron RBD domain by COVOX-222. Mutation sites correspond to the Omicron RBD are labeled in red (1-15) and are listed in Figure 4B. Table summarizing the protein structure used for primary models for analyzing the molecular dynamics of COVOX-222 interaction with Omicron RBD. The table also includes amino acid positions, the corresponding -KL divergence value denoting atomic fluctuations dampening, D value and the corresponding level of significance for the KL divergence values.

**Supplemental Figure 2:**
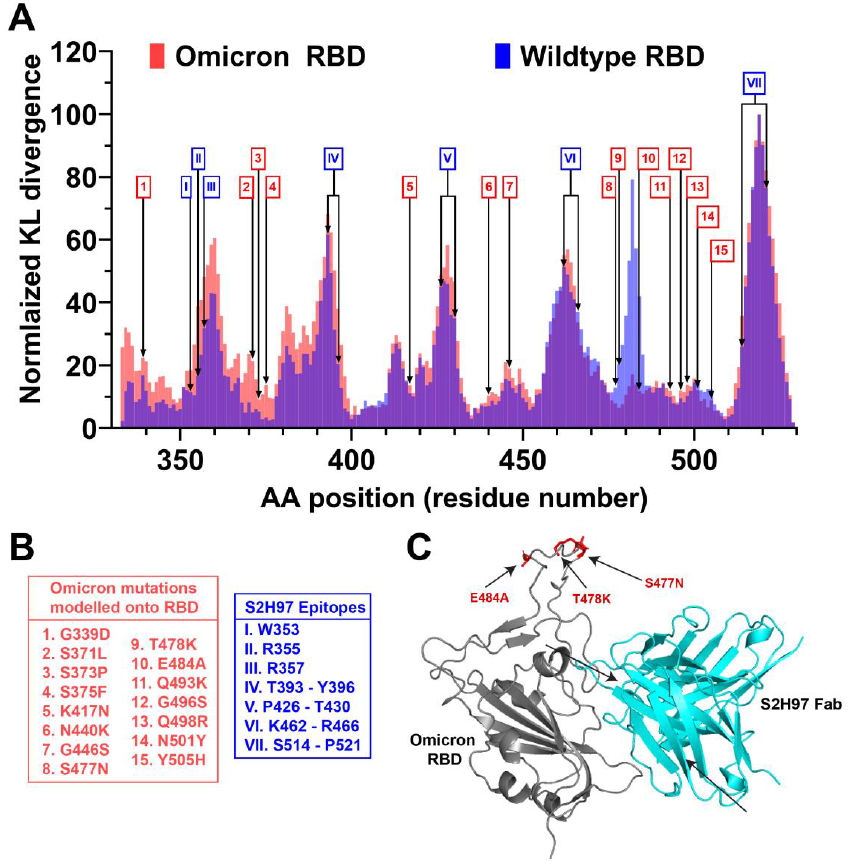
MD simulations with the Omicron variant reveal sites that promote similiar binding affinity to hACE2. (A) Sequence positional plotting of the normalized dampening of atom motions on the Omicron RBD (red) and the wildtype RBD (blue) by monoclonal antibody S2H97.. The omicron RBD mutations are labeled in red (1-15). The sites corresponding to the S2H97 epitope on the wildtype RBD on labeled in blue (I – VII). The amino acid residues that correspond to the numbers and roman numerals are listed in (B). In the omicron RBD, we see alteration in atomic fluctuation peaks at several of the epitope sites.. (B) List of omicron mutations modeled onto RBD, and the list of S2H97 epitopes. (C) Crystal structure of S2H97 Fab (cyan) superimposed onto the structure of the Omicron RBD (grey) (PDB 7M7W). Some of the amino acid residues that correspond to the S2H97 epitopes, show an increase in atomic fluctuation dampening, are shown in blue. Several of the amino acid residue that correspond to omicron mutations with increased atomic fluctuation dampening, which are sites of ACE2 interactions, are shown in red. NOTE: Major changes occurring E4848A, T478K, S477N.

