## Supplementary material for "Identifying protein sites contributing to vaccine escape via statistical comparisons of short-term molecular dynamics simulations": data_extraction.ipynb

```
# %% [code]
import pandas as pd
data = pd.read_csv("DROIDSfluctuationAVGchain.txt")
data2 = pd.read_csv("adjustKStests.txt")

# %% [code]
new = data["pos_ref\tres_ref\tres_query\tflux_ref_avg\tflux_query_avg\tdelta_flux\tabs_delta_flux\tKLdivergence\tchain"].str.split("\t",9,expand = True)
new2 = data2["posAA\trefAA\tqueryAA\tDval\tpval\tsignif"].str.split("\t",5,expand=True)

# %% [code]
data["pos_ref"]=new[0]
data["res_ref"]=new[1]
data["flux_ref_avg"]=new[3]
data["flux_query_avg"]=new[4]
data["KLdivergence"]=new[7]
data=data.drop(["pos_ref\tres_ref\tres_query\tflux_ref_avg\tflux_query_avg\tdelta_flux\tabs_delta_flux\tKLdivergence\tchain"],axis=1)
data["Dvalue"]=new2[3]
data["pvalue"]=new2[4]
data["significance"]=new2[5]

# %% [code]
aa_list = data["res_ref"].value_counts().sort_index(ascending=True).index.to_list()

# %% [code]
data.head()

# %% [code]
repl_value = {"<0.1alpha":"p < 0.1", "<0.5alpha":"p < 0.5", "ns" : "No significance", "<1alpha":"p < 1"}
data = data.replace({"significance":repl_value})
sig_list = data["significance"].value_counts().sort_index(ascending=True).index.to_list()

# %% [code]
for aa in aa_list:
    data[aa]=data.loc[data["res_ref"]==aa,"KLdivergence"]
for value in sig_list:
    data[value]=data.loc[data["significance"]==value,"Dvalue"]
data = data.fillna("")

# %% [code]
data.to_csv("data_plot.csv",index=False)

# %% [code]
```
