## Supplementary figures and images for "Identifying protein sites contributing to vaccine escape via statistical comparisons of short-term molecular dynamics simulations"

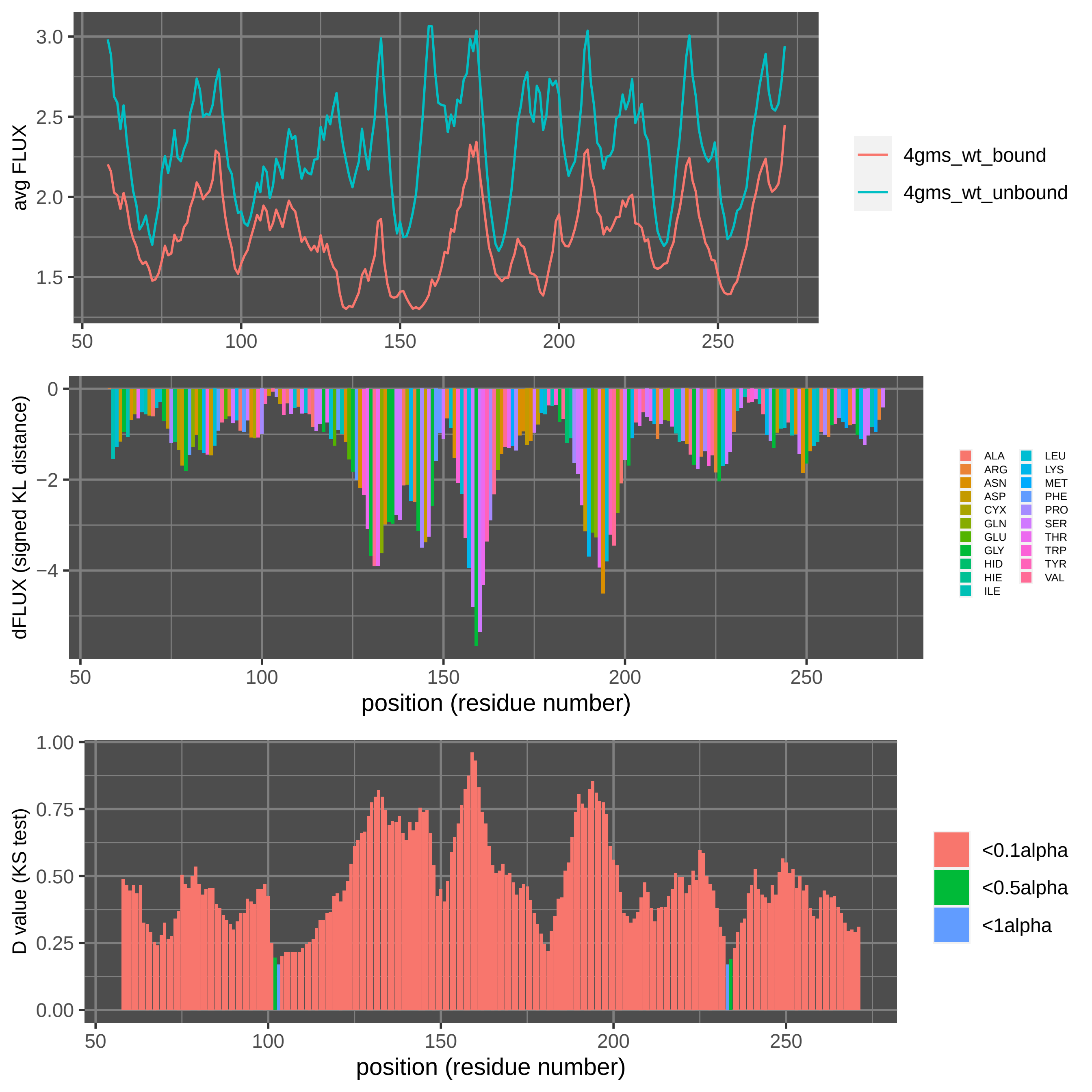

Supplement: Supplemental Data [file 471374_file09.zip › data_RajendranFerranBabbitt_2022/Figure 2A [influenza HA head + S139_1)/DROIDSplot_all.pdf]

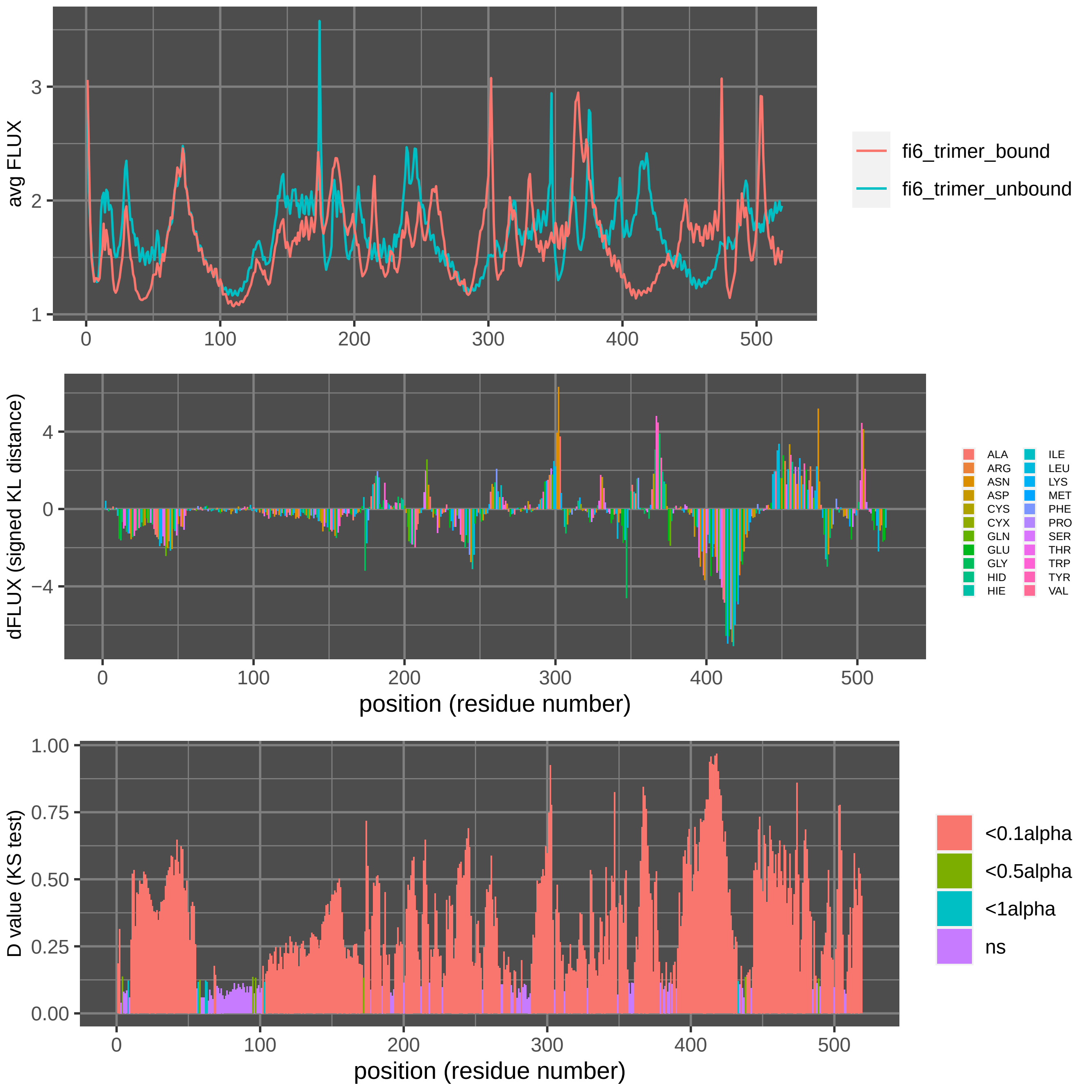

Supplement: Supplemental Data [file 471374_file09.zip › data_RajendranFerranBabbitt_2022/Figure 2C [influenza HA stalk + FI6v3]/DROIDSplot_all.pdf]

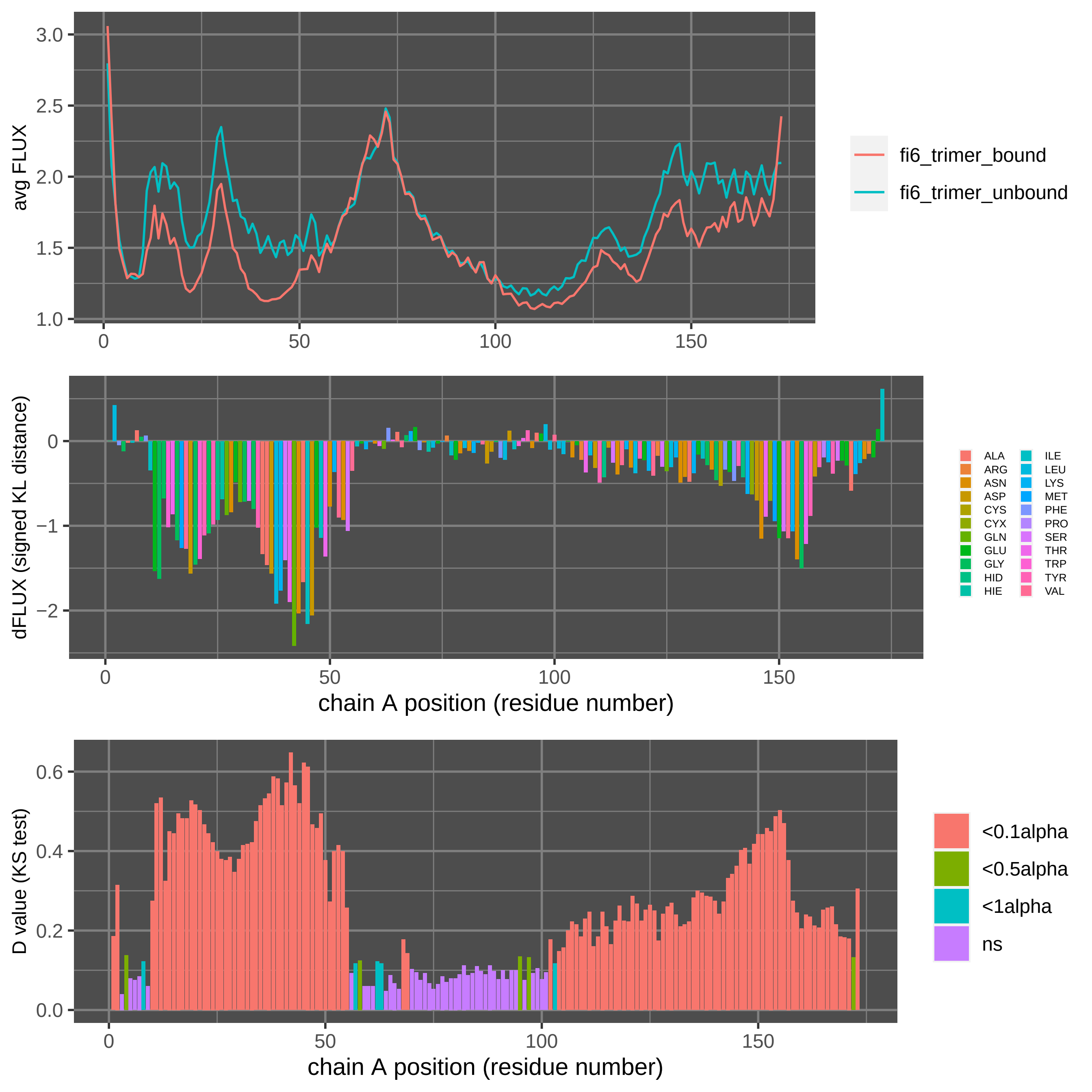

Supplement: Supplemental Data [file 471374_file09.zip › data_RajendranFerranBabbitt_2022/Figure 2C [influenza HA stalk + FI6v3]/DROIDS_results_fi6_trimer_bound_fi6_trimer_unbound_flux_0.010_A/DROIDSplot_A.pdf]

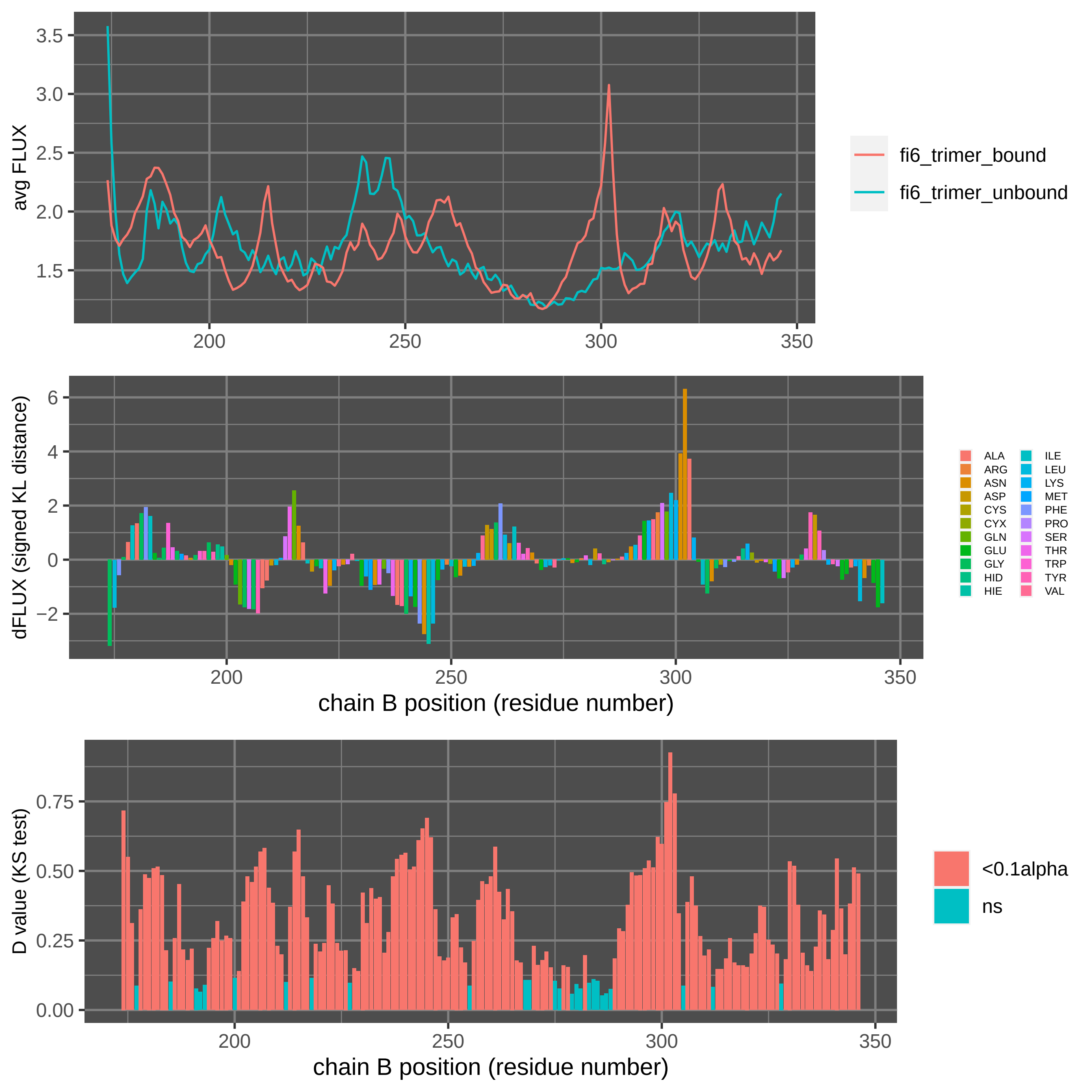

Supplement: Supplemental Data [file 471374_file09.zip › data_RajendranFerranBabbitt_2022/Figure 2C [influenza HA stalk + FI6v3]/DROIDS_results_fi6_trimer_bound_fi6_trimer_unbound_flux_0.010_B/DROIDSplot_B.pdf]

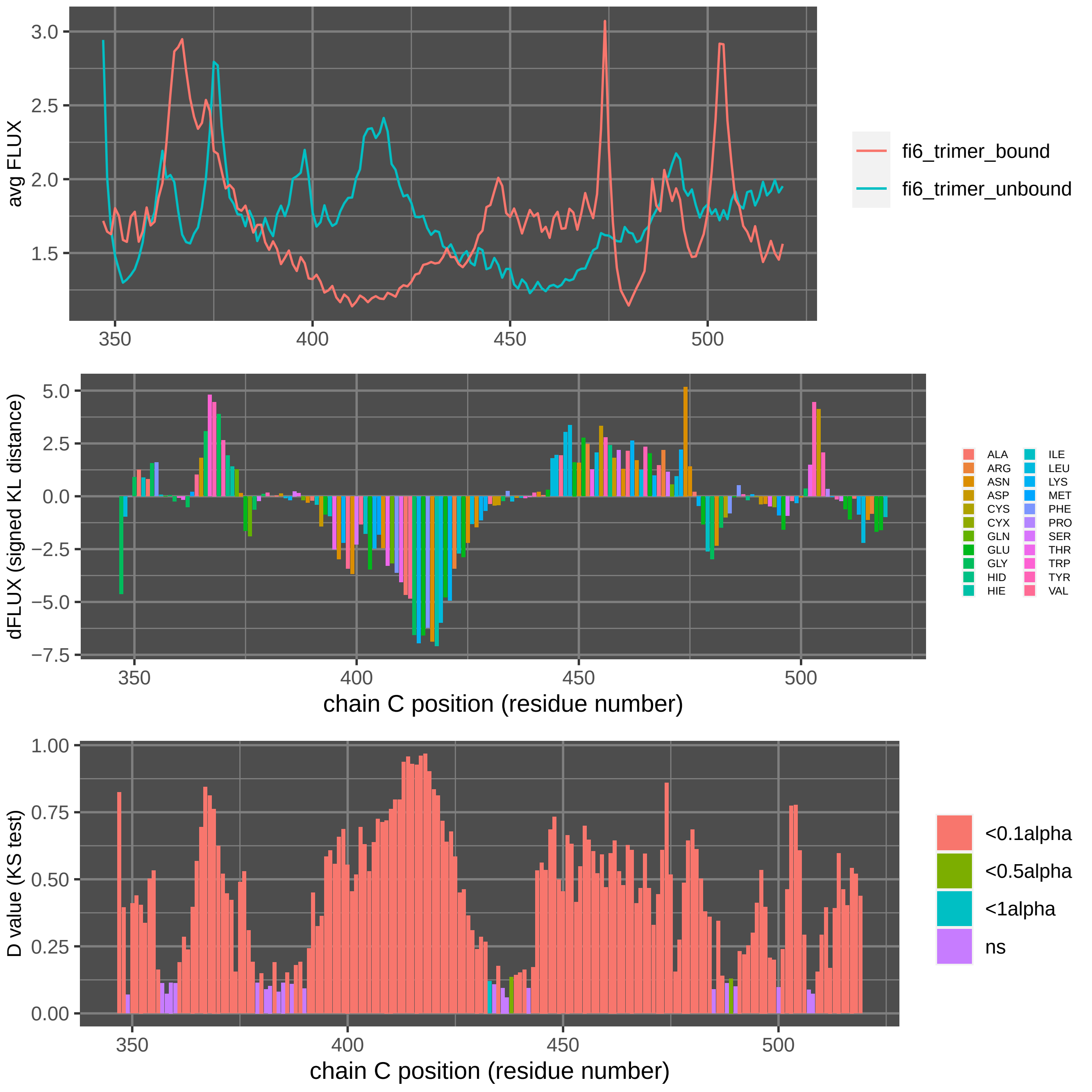

Supplement: Supplemental Data [file 471374_file09.zip › data_RajendranFerranBabbitt_2022/Figure 2C [influenza HA stalk + FI6v3]/DROIDS_results_fi6_trimer_bound_fi6_trimer_unbound_flux_0.010_C/DROIDSplot_C.pdf]

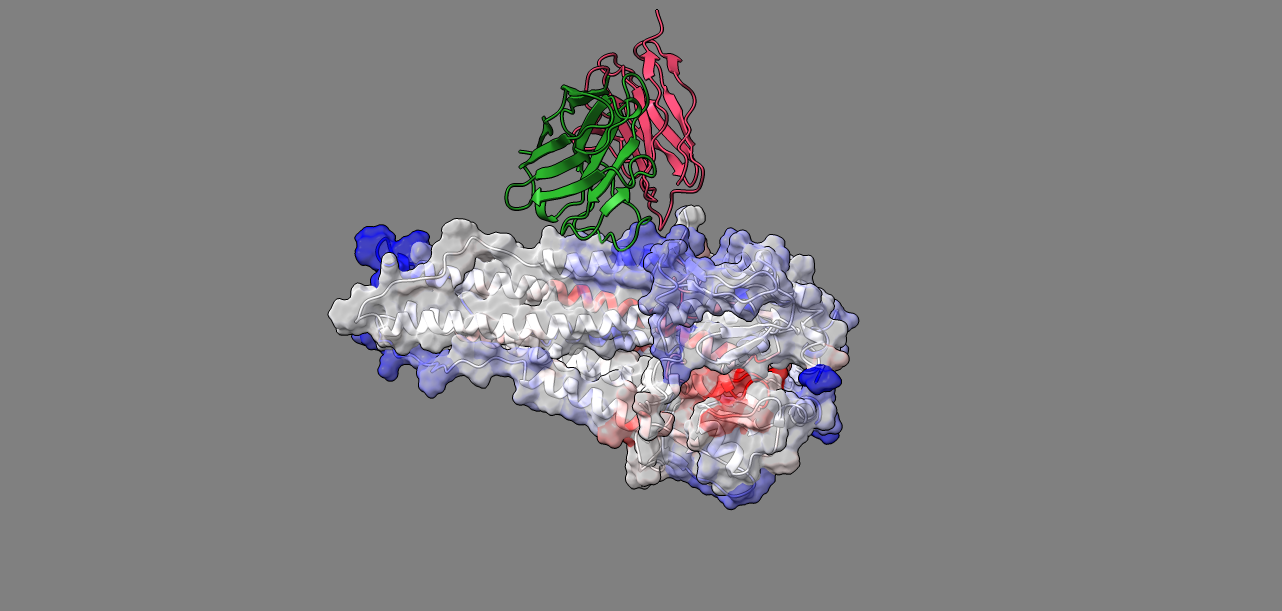

Supplement: Supplemental Data [file 471374_file09.zip › data_RajendranFerranBabbitt_2022/Figure 2C [influenza HA stalk + FI6v3]/image1.png]

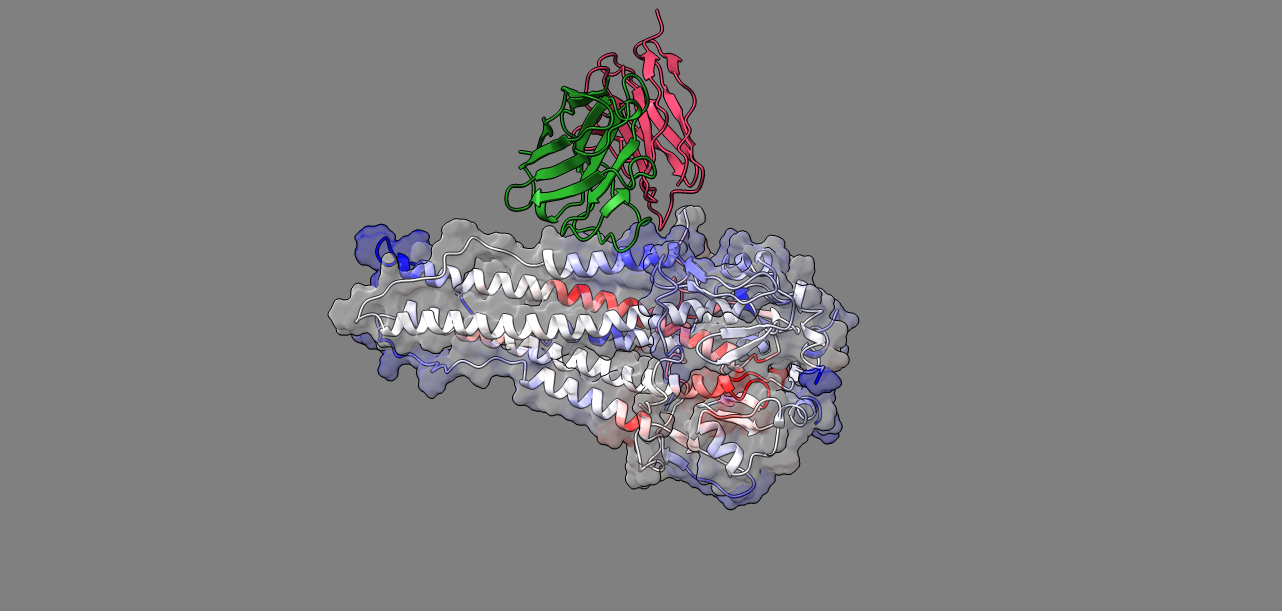

Supplement: Supplemental Data [file 471374_file09.zip › data_RajendranFerranBabbitt_2022/Figure 2C [influenza HA stalk + FI6v3]/image3.png]

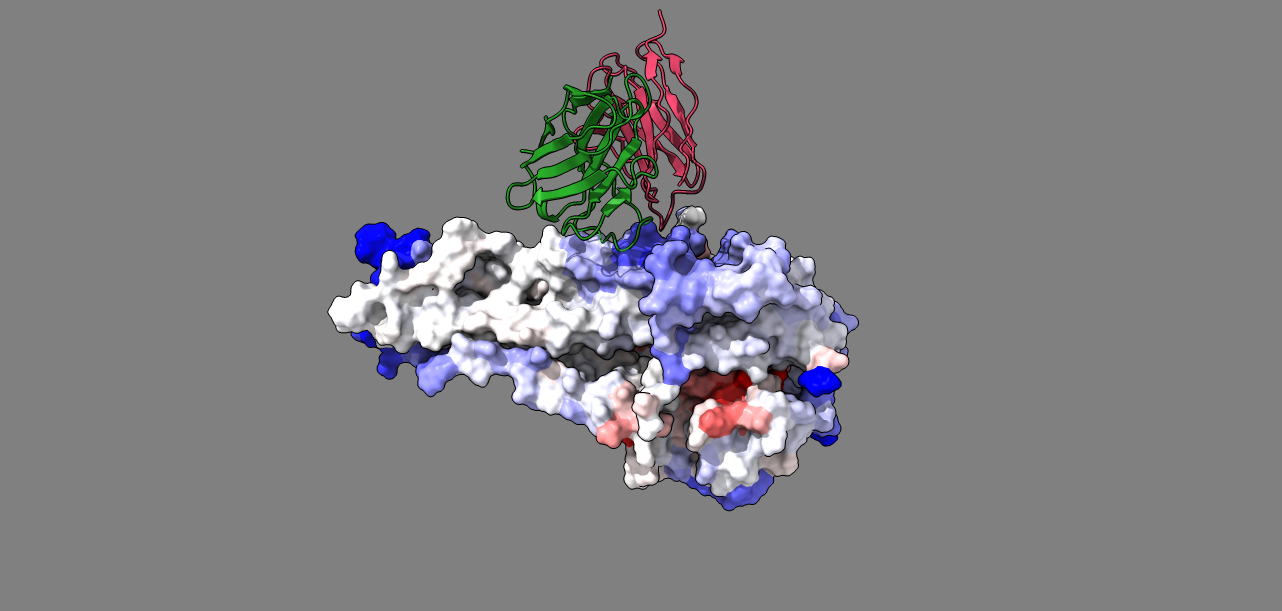

Supplement: Supplemental Data [file 471374_file09.zip › data_RajendranFerranBabbitt_2022/Figure 2C [influenza HA stalk + FI6v3]/image4.png]

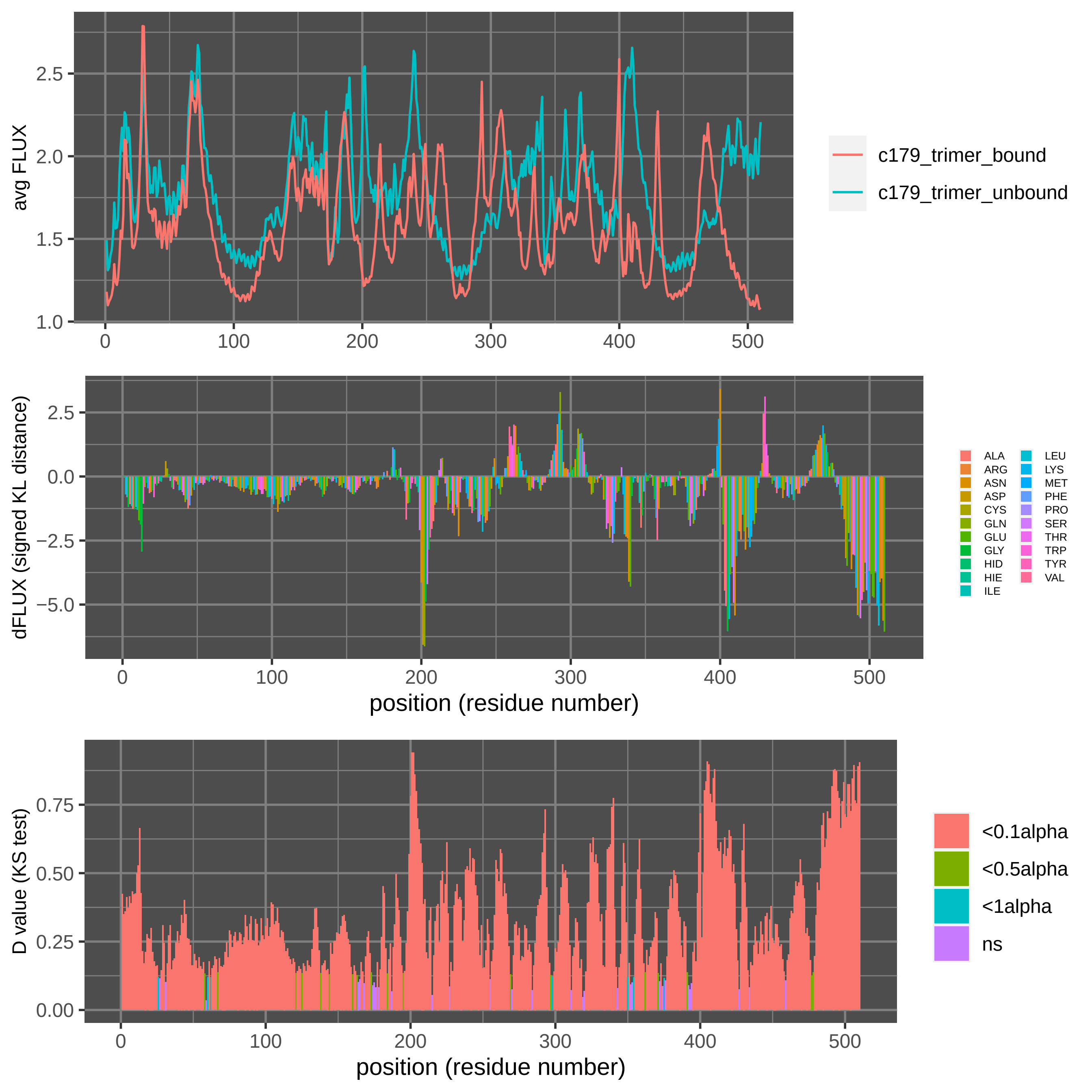

Supplement: Supplemental Data [file 471374_file09.zip › data_RajendranFerranBabbitt_2022/Figure 2D [influenza HA stalk + C179]/DROIDS_results_c179_trimer_bound_c179_trimer_unbound_flux_0.010/DROIDSplot_all.pdf]

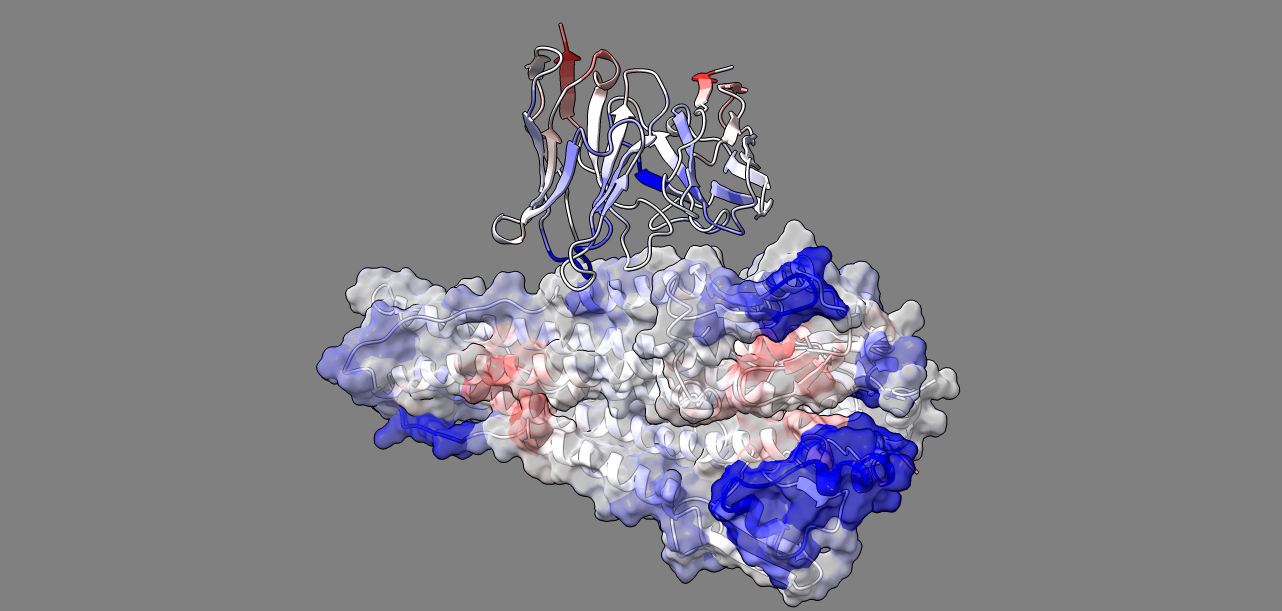

Supplement: Supplemental Data [file 471374_file09.zip › data_RajendranFerranBabbitt_2022/Figure 2D [influenza HA stalk + C179]/DROIDS_results_c179_trimer_bound_c179_trimer_unbound_flux_0.010/image5.png]

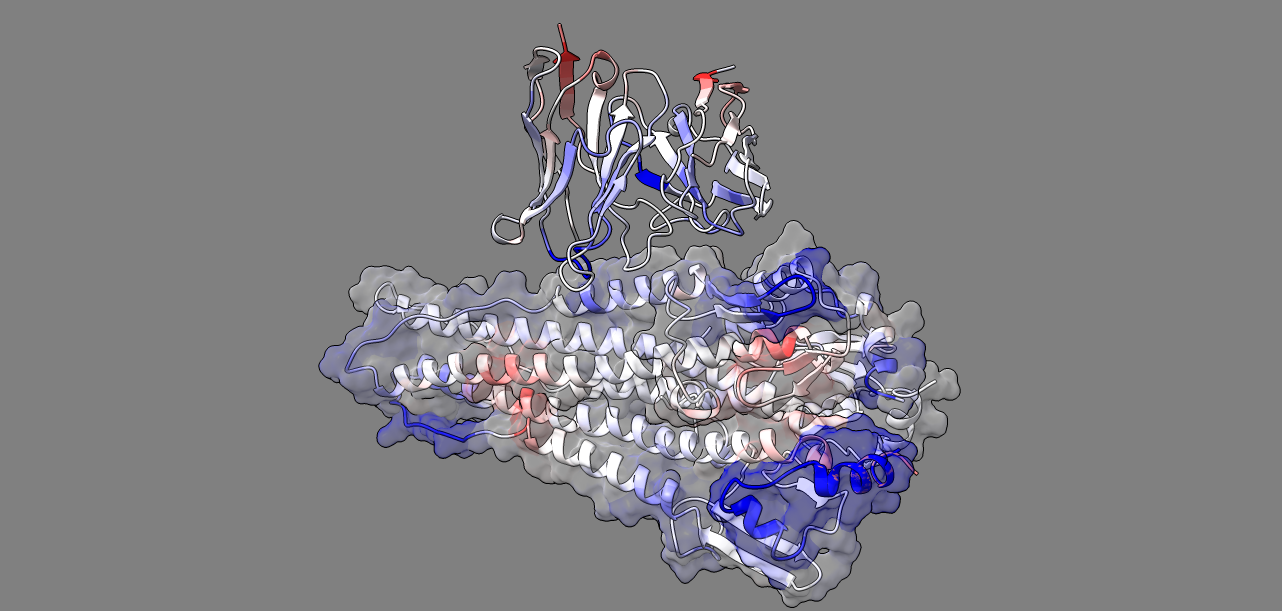

Supplement: Supplemental Data [file 471374_file09.zip › data_RajendranFerranBabbitt_2022/Figure 2D [influenza HA stalk + C179]/DROIDS_results_c179_trimer_bound_c179_trimer_unbound_flux_0.010/image6.png]

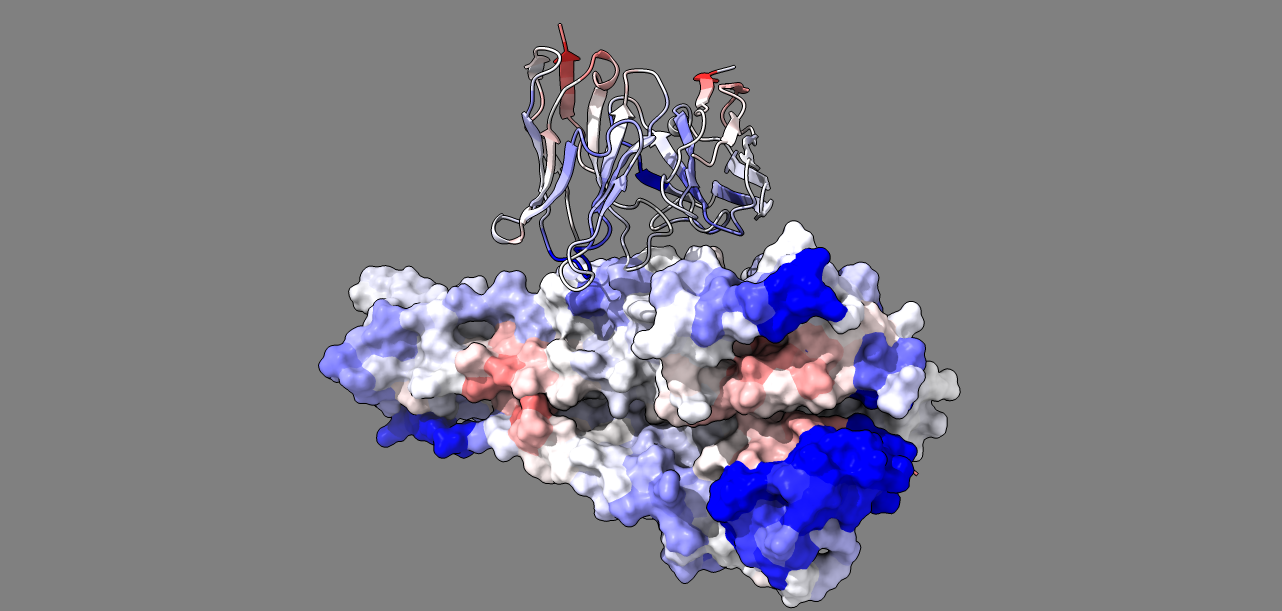

Supplement: Supplemental Data [file 471374_file09.zip › data_RajendranFerranBabbitt_2022/Figure 2D [influenza HA stalk + C179]/DROIDS_results_c179_trimer_bound_c179_trimer_unbound_flux_0.010/image7.png]

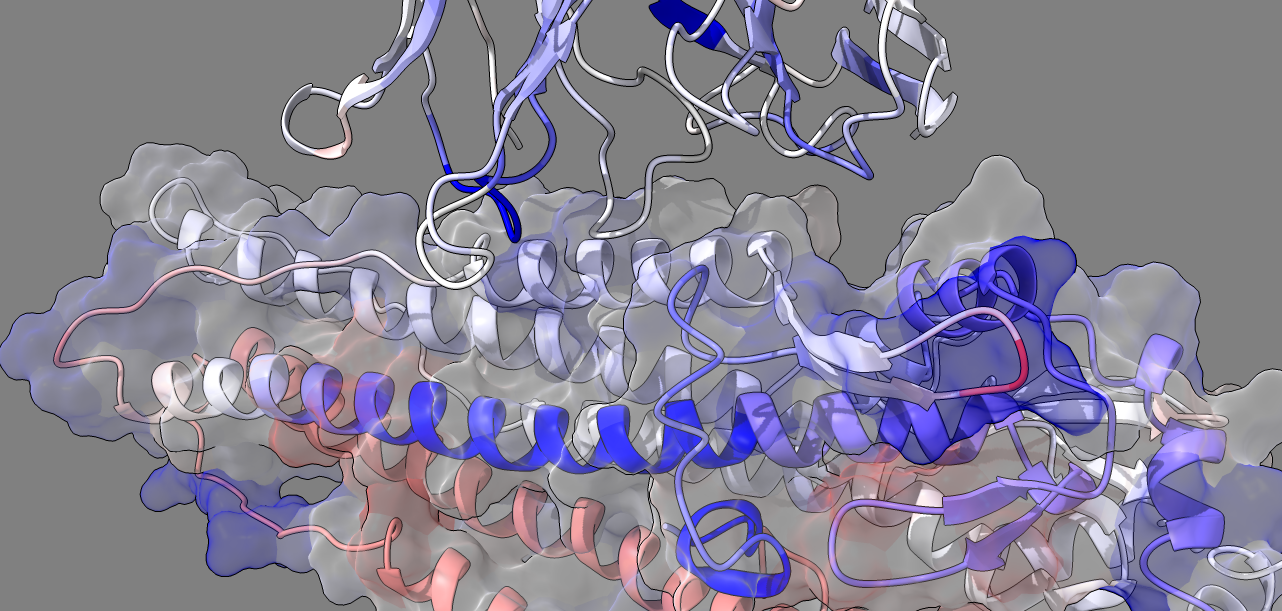

Supplement: Supplemental Data [file 471374_file09.zip › data_RajendranFerranBabbitt_2022/Figure 2D [influenza HA stalk + C179]/DROIDS_results_c179_trimer_bound_c179_trimer_unbound_flux_0.010/image8.png]

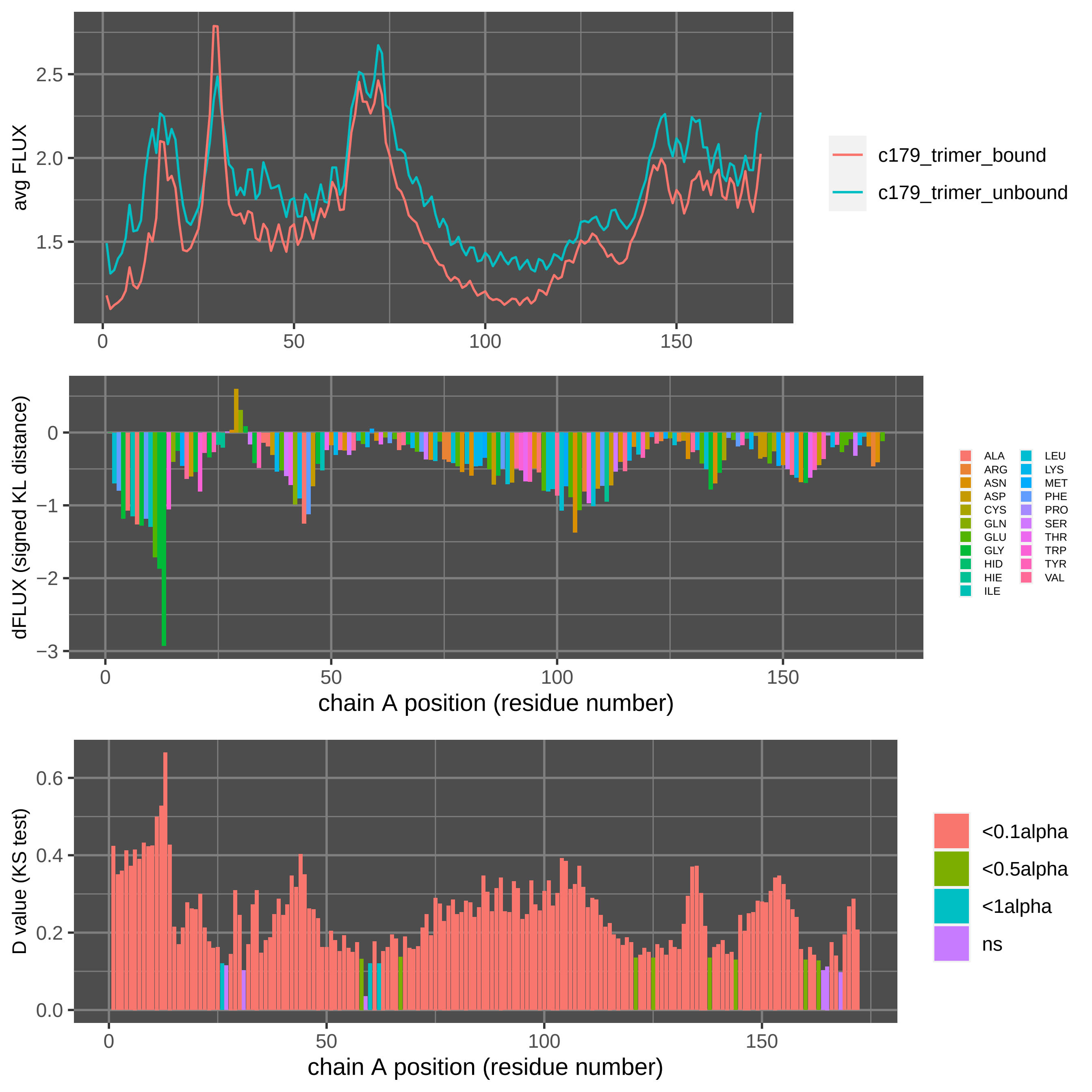

Supplement: Supplemental Data [file 471374_file09.zip › data_RajendranFerranBabbitt_2022/Figure 2D [influenza HA stalk + C179]/DROIDS_results_c179_trimer_bound_c179_trimer_unbound_flux_0.010_A/DROIDSplot_A.pdf]

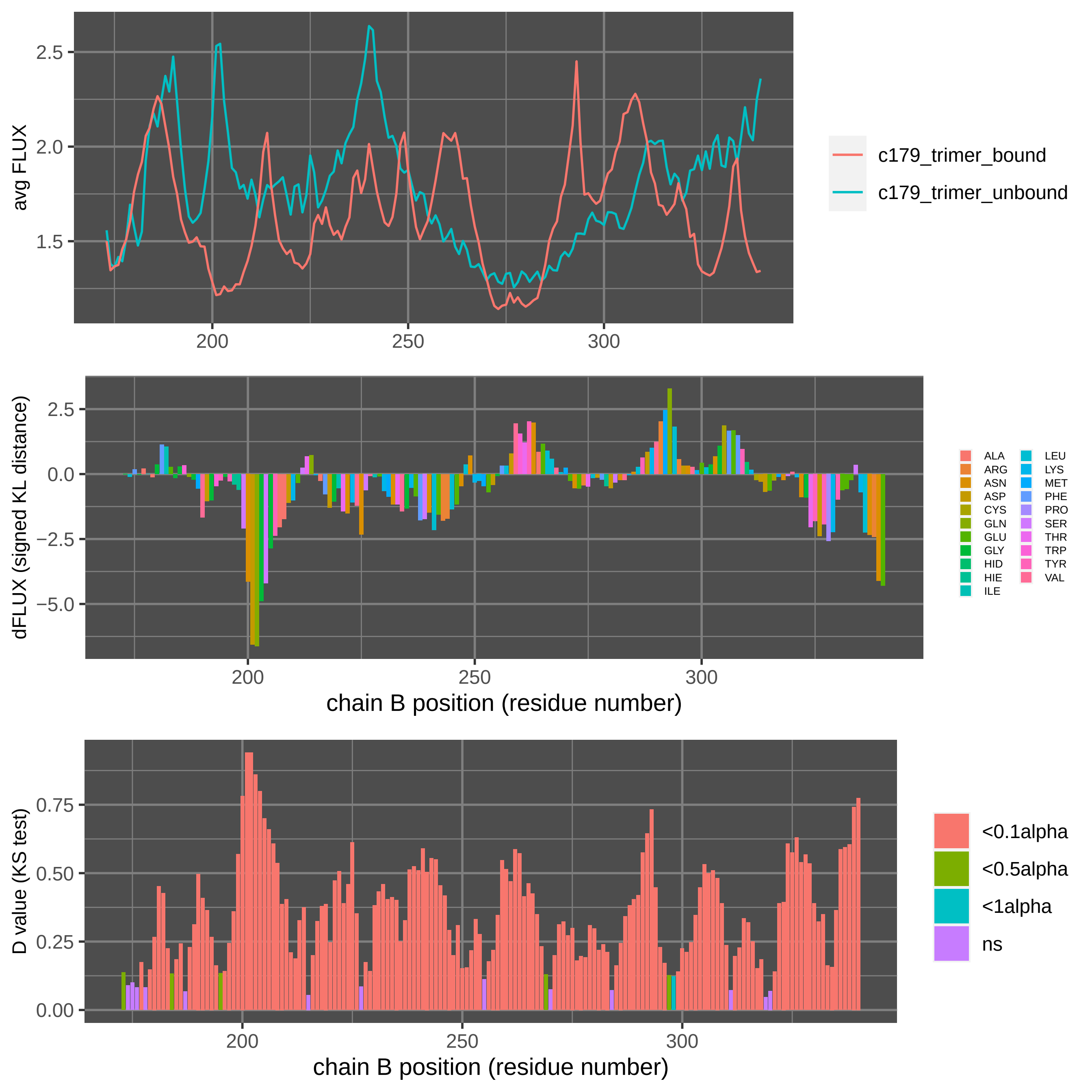

Supplement: Supplemental Data [file 471374_file09.zip › data_RajendranFerranBabbitt_2022/Figure 2D [influenza HA stalk + C179]/DROIDS_results_c179_trimer_bound_c179_trimer_unbound_flux_0.010_B/DROIDSplot_B.pdf]

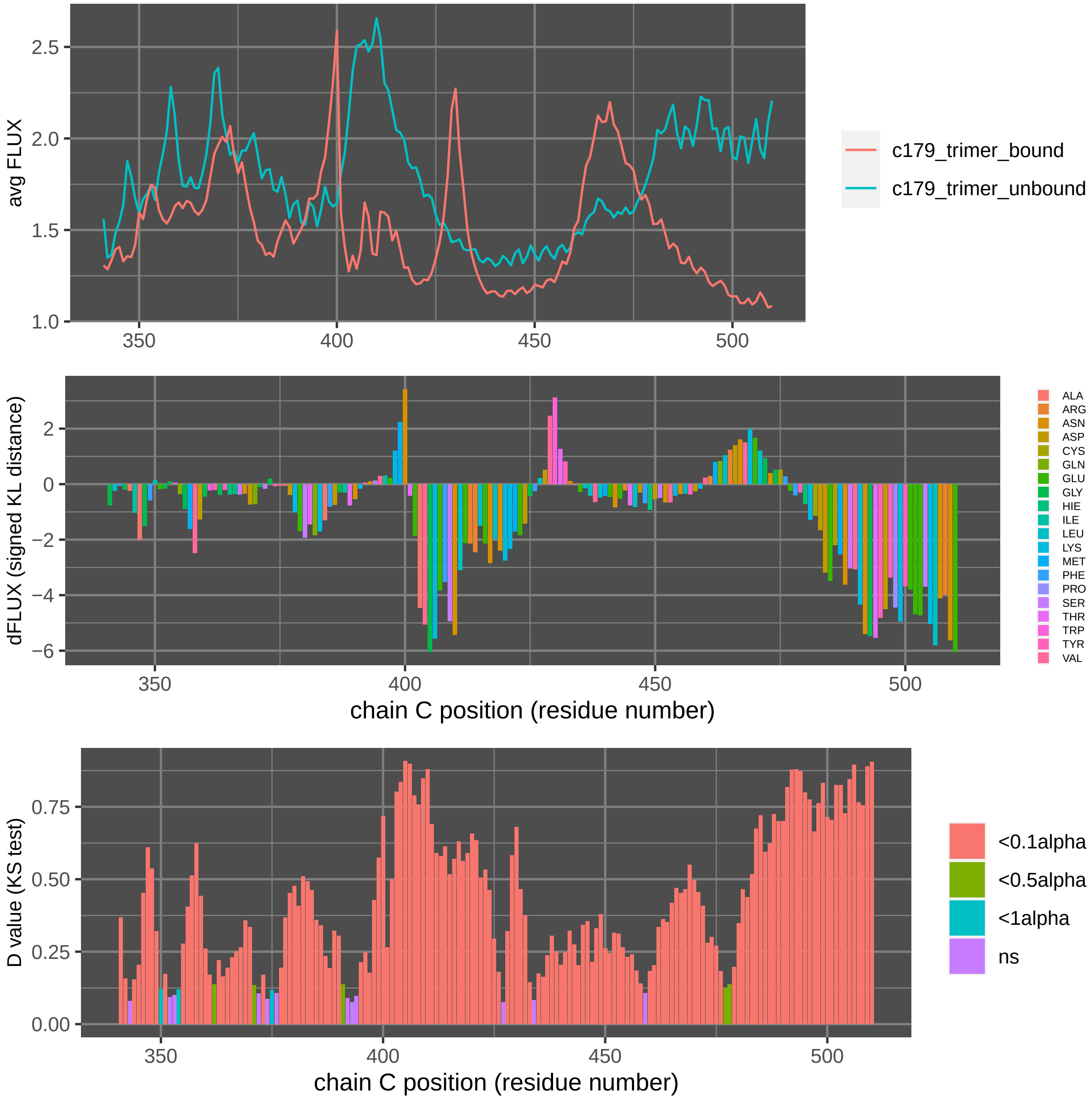

Supplement: Supplemental Data [file 471374_file09.zip › data_RajendranFerranBabbitt_2022/Figure 2D [influenza HA stalk + C179]/DROIDS_results_c179_trimer_bound_c179_trimer_unbound_flux_0.010_C/DROIDSplot_C.pdf]

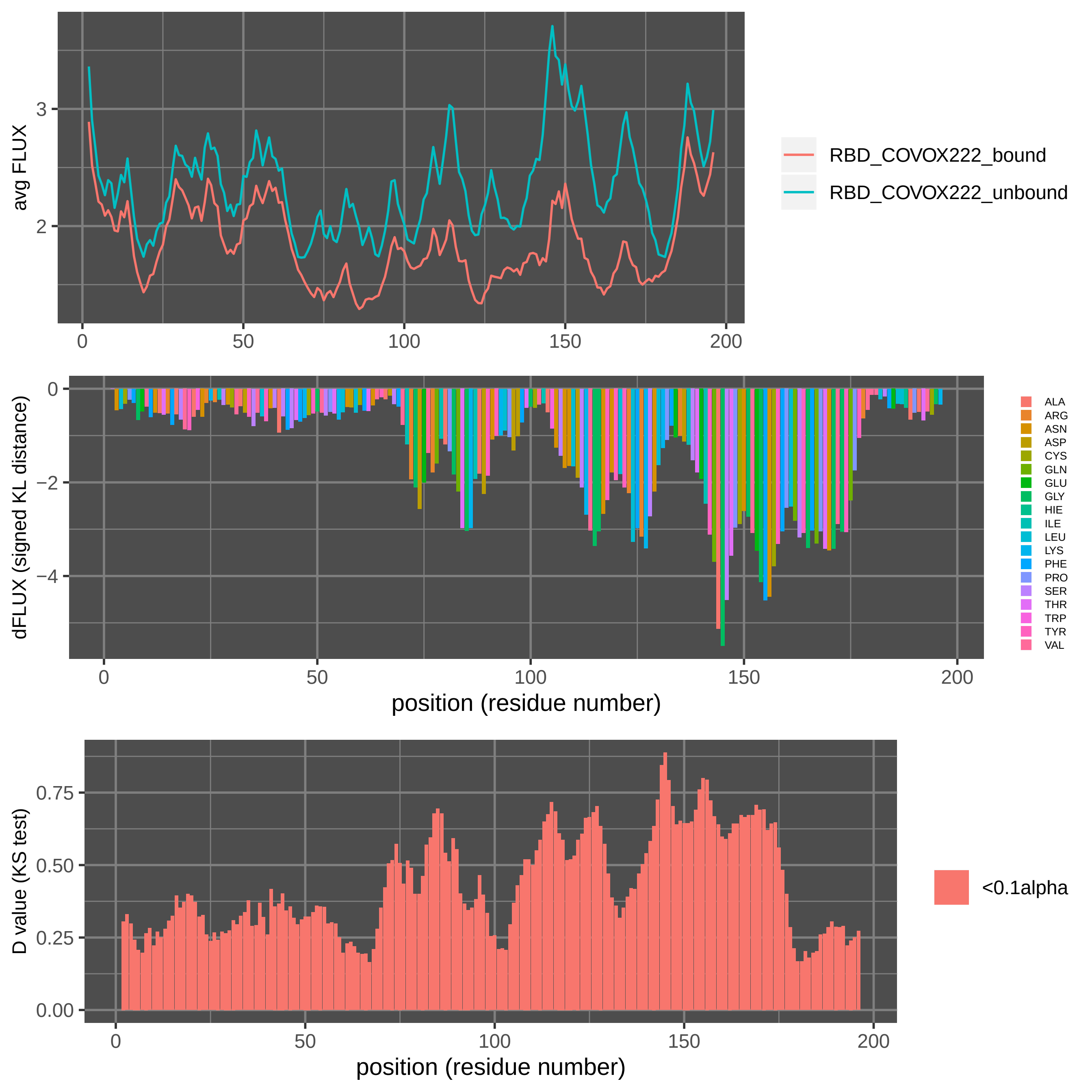

Supplement: Supplemental Data [file 471374_file09.zip › data_RajendranFerranBabbitt_2022/Figure 3B[Sars-CoV-2 RBD + COVOX-222]/DROIDSplot_all.pdf]

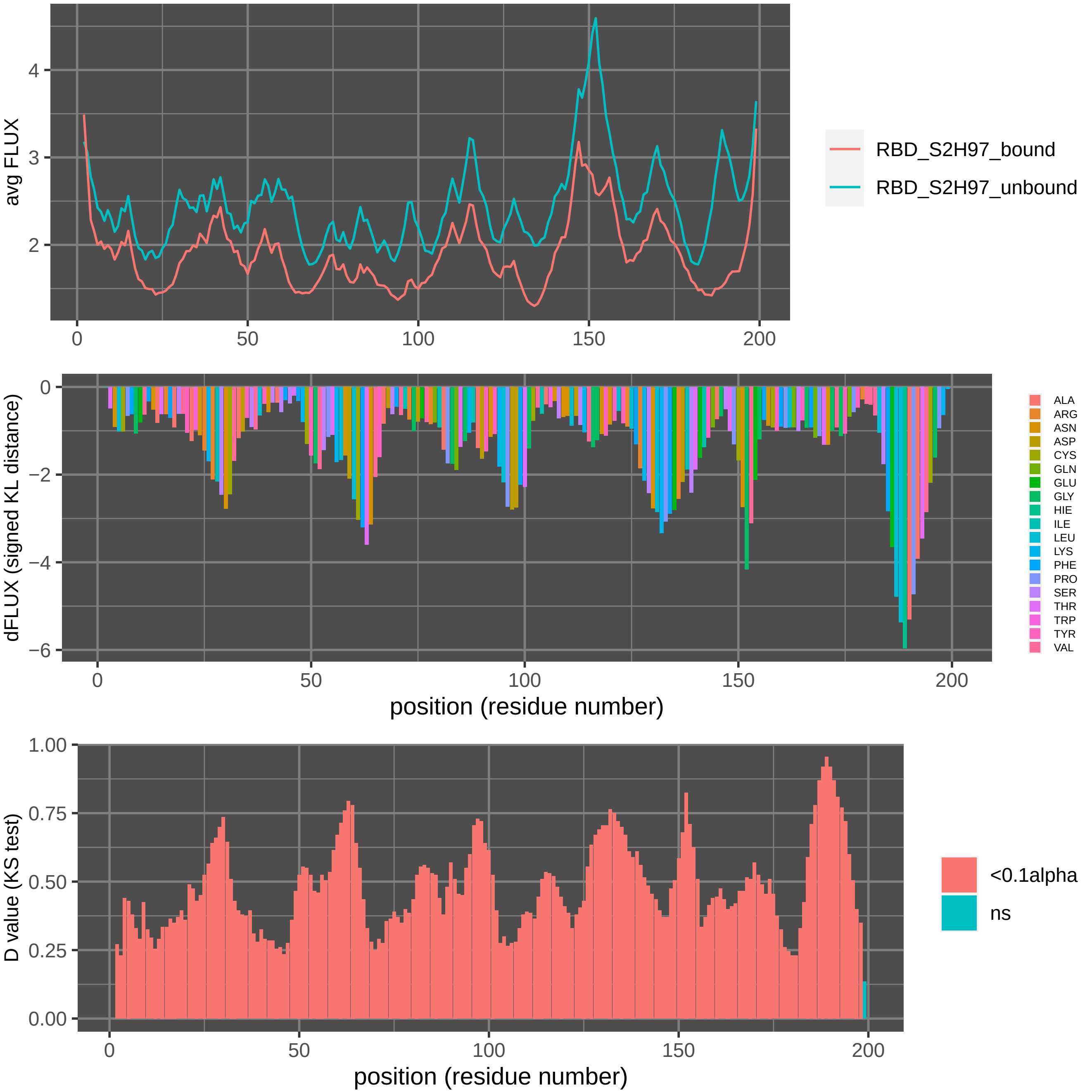

Supplement: Supplemental Data [file 471374_file09.zip › data_RajendranFerranBabbitt_2022/Figure 3C[Sars-CoV-2 RBD + S2H97]/DROIDSplot_.pdf]

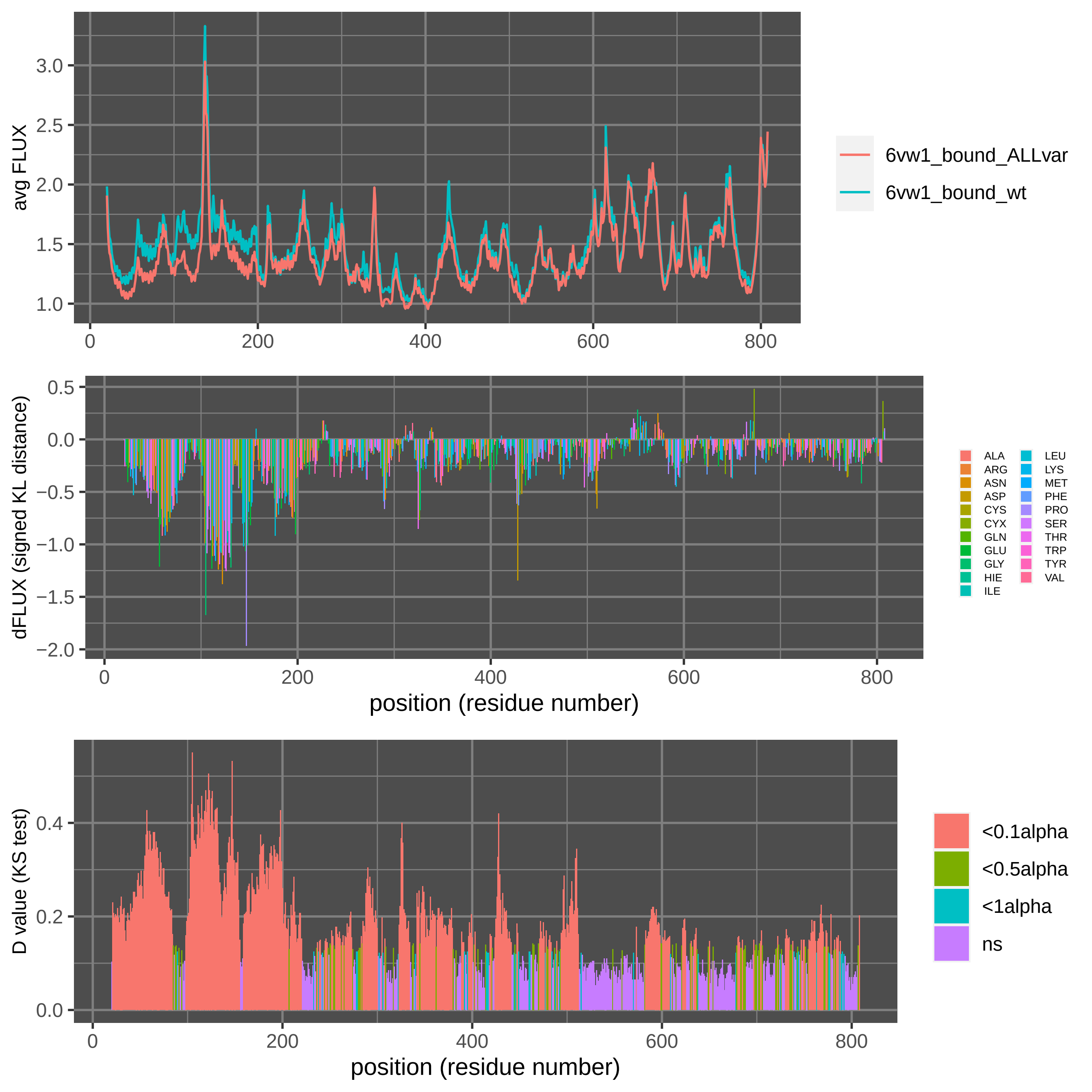

Supplement: Supplemental Data [file 471374_file09.zip › data_RajendranFerranBabbitt_2022/Figure 3D,3E [Sars-CoV-2 RBD WT_Mut(voc) + ACE2]/all_variants_ACE2bound/DROIDS_results_6vw1_bound_ALLvar_6vw1_bound_wt_flux_0.010/DROIDSplot_all.pdf]

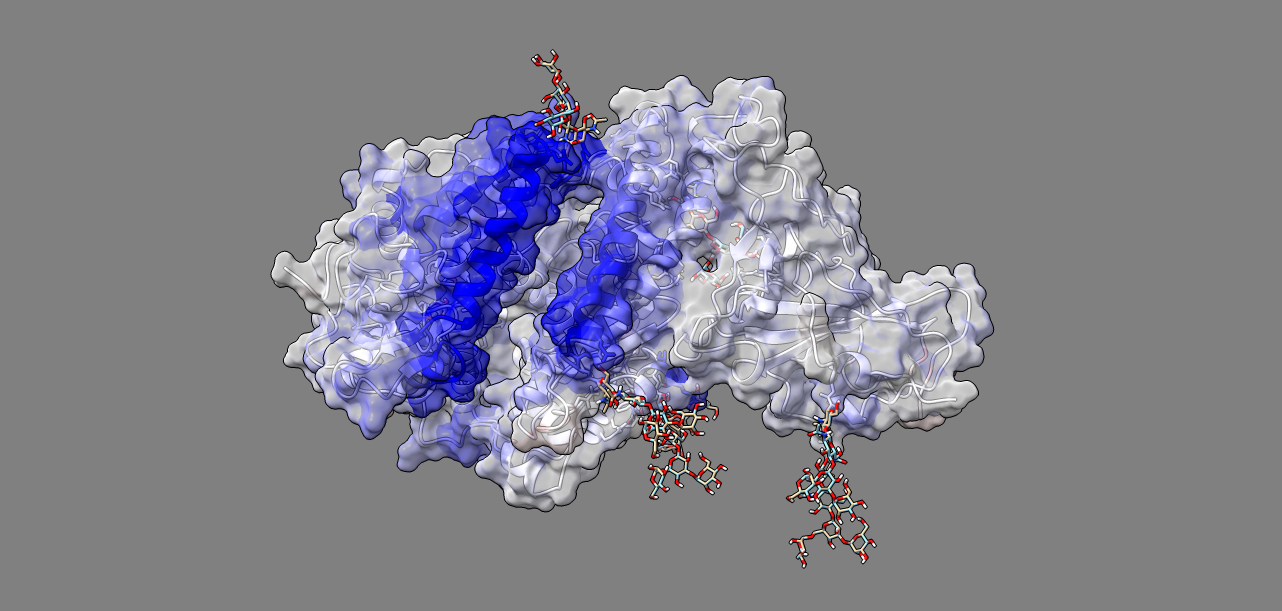

Supplement: Supplemental Data [file 471374_file09.zip › data_RajendranFerranBabbitt_2022/Figure 3D,3E [Sars-CoV-2 RBD WT_Mut(voc) + ACE2]/all_variants_ACE2bound/DROIDS_results_6vw1_bound_ALLvar_6vw1_bound_wt_flux_0.010/image1.png]

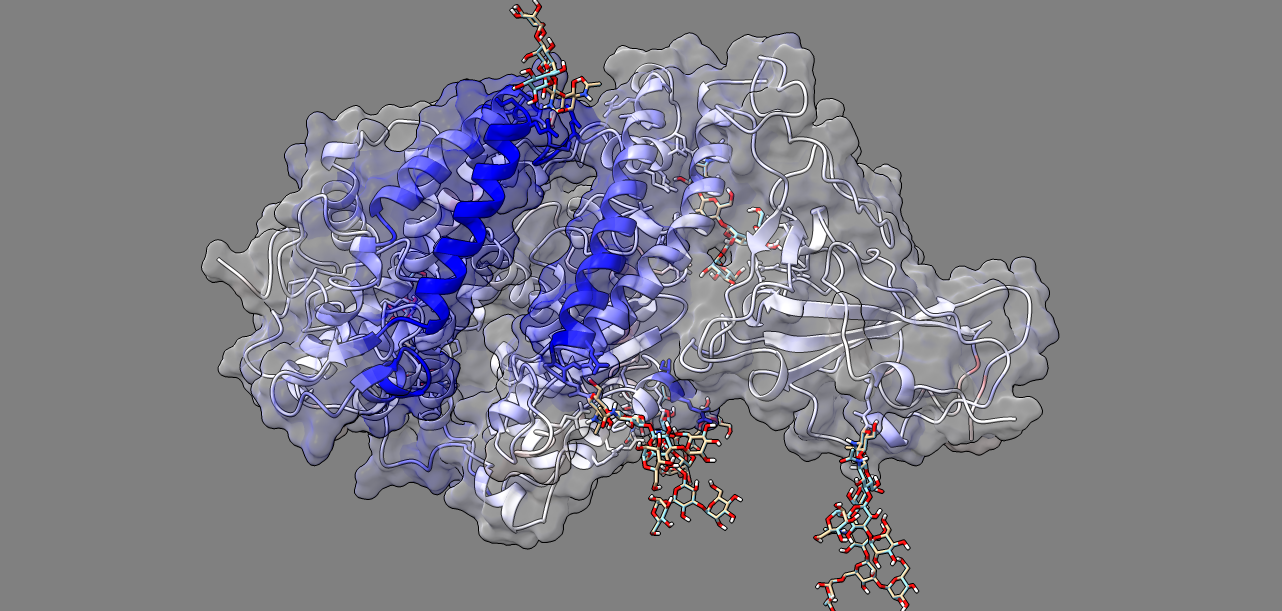

Supplement: Supplemental Data [file 471374_file09.zip › data_RajendranFerranBabbitt_2022/Figure 3D,3E [Sars-CoV-2 RBD WT_Mut(voc) + ACE2]/all_variants_ACE2bound/DROIDS_results_6vw1_bound_ALLvar_6vw1_bound_wt_flux_0.010/image2.png]

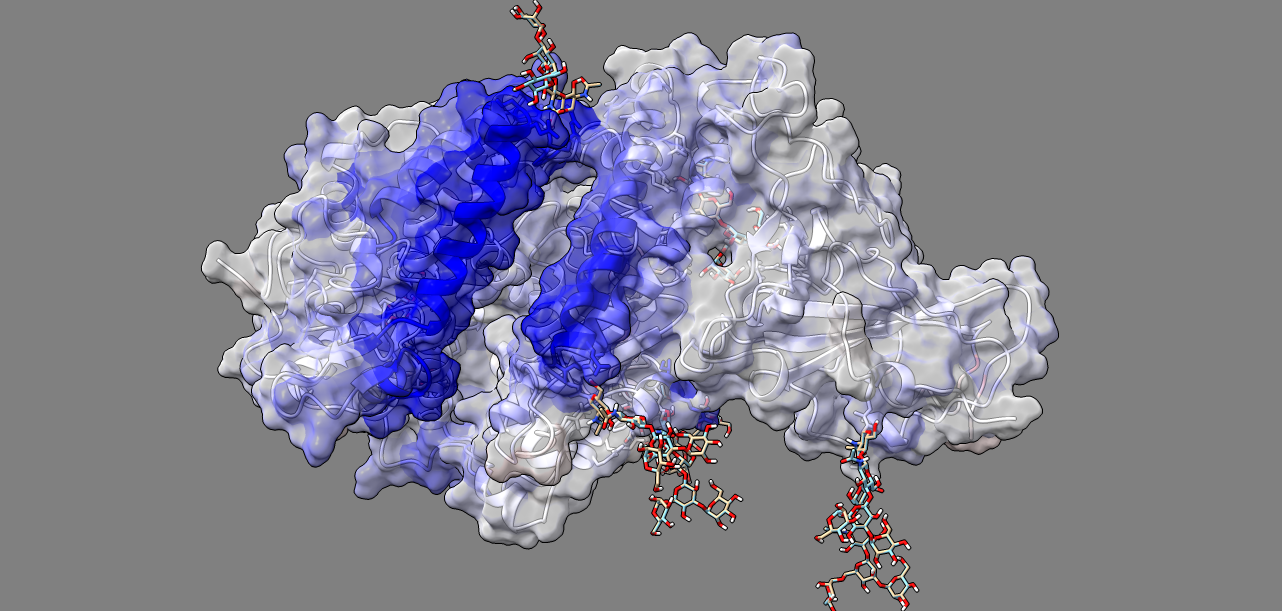

Supplement: Supplemental Data [file 471374_file09.zip › data_RajendranFerranBabbitt_2022/Figure 3D,3E [Sars-CoV-2 RBD WT_Mut(voc) + ACE2]/all_variants_ACE2bound/DROIDS_results_6vw1_bound_ALLvar_6vw1_bound_wt_flux_0.010/image3.png]

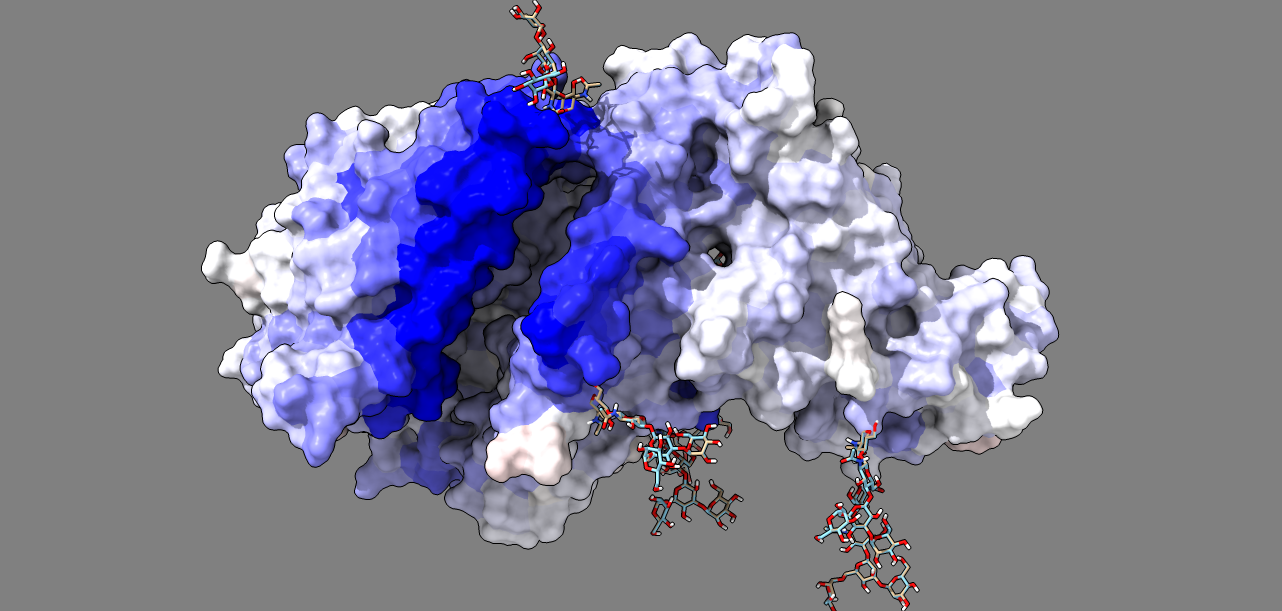

Supplement: Supplemental Data [file 471374_file09.zip › data_RajendranFerranBabbitt_2022/Figure 3D,3E [Sars-CoV-2 RBD WT_Mut(voc) + ACE2]/all_variants_ACE2bound/DROIDS_results_6vw1_bound_ALLvar_6vw1_bound_wt_flux_0.010/image4.png]

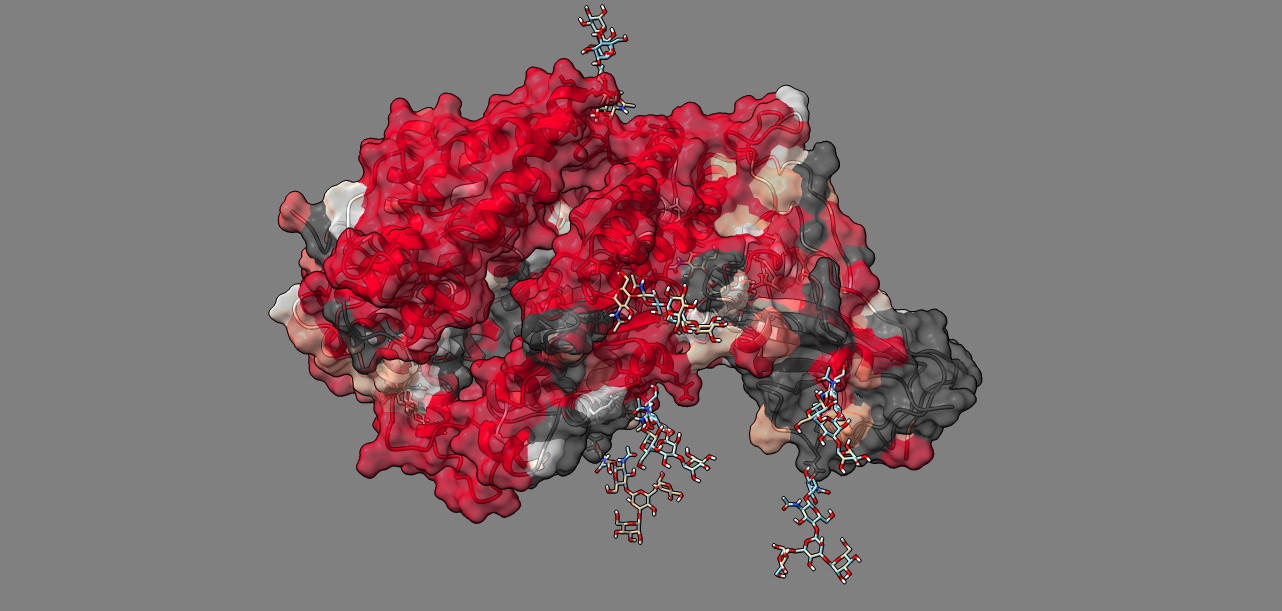

Supplement: Supplemental Data [file 471374_file09.zip › data_RajendranFerranBabbitt_2022/Figure 3D,3E [Sars-CoV-2 RBD WT_Mut(voc) + ACE2]/all_variants_ACE2bound/DROIDS_results_6vw1_bound_ALLvar_6vw1_bound_wt_flux_0.010/image5.png]

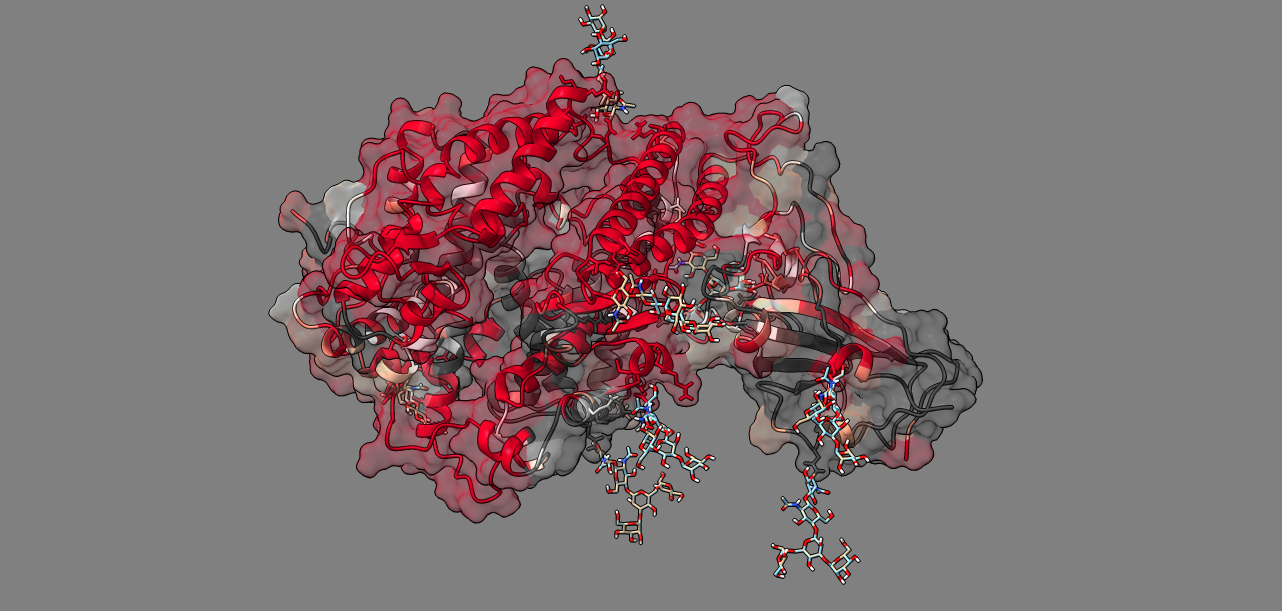

Supplement: Supplemental Data [file 471374_file09.zip › data_RajendranFerranBabbitt_2022/Figure 3D,3E [Sars-CoV-2 RBD WT_Mut(voc) + ACE2]/all_variants_ACE2bound/DROIDS_results_6vw1_bound_ALLvar_6vw1_bound_wt_flux_0.010/image6.png]

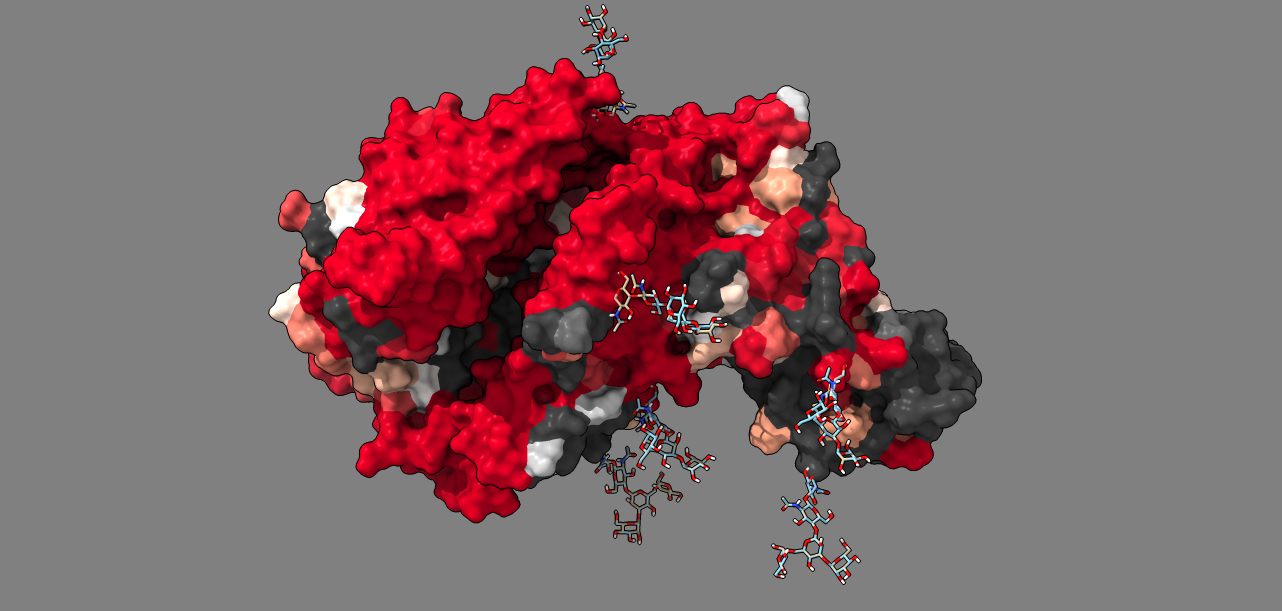

Supplement: Supplemental Data [file 471374_file09.zip › data_RajendranFerranBabbitt_2022/Figure 3D,3E [Sars-CoV-2 RBD WT_Mut(voc) + ACE2]/all_variants_ACE2bound/DROIDS_results_6vw1_bound_ALLvar_6vw1_bound_wt_flux_0.010/image7.png]

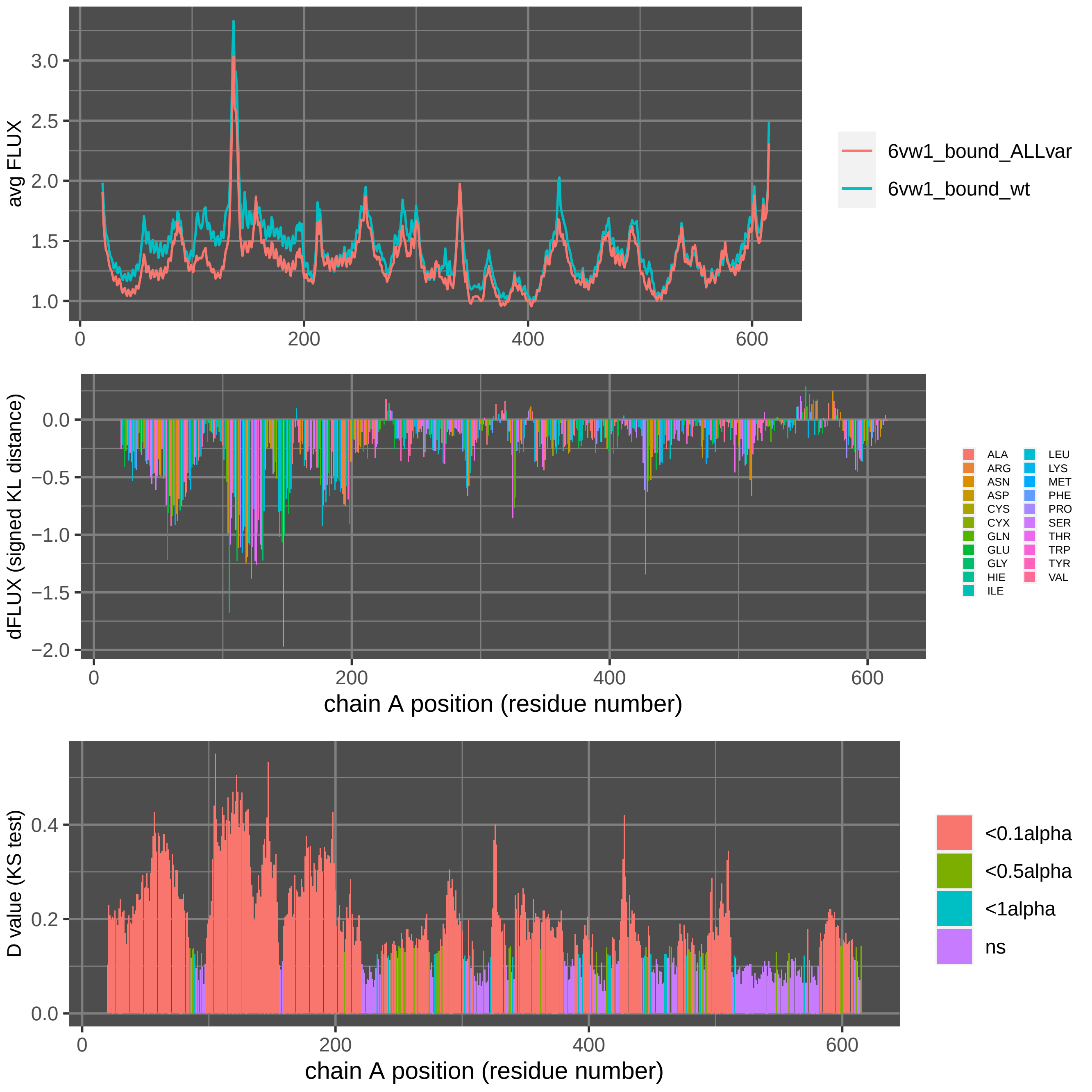

Supplement: Supplemental Data [file 471374_file09.zip › data_RajendranFerranBabbitt_2022/Figure 3D,3E [Sars-CoV-2 RBD WT_Mut(voc) + ACE2]/all_variants_ACE2bound/DROIDS_results_6vw1_bound_ALLvar_6vw1_bound_wt_flux_0.010_A/DROIDSplot_A.pdf]

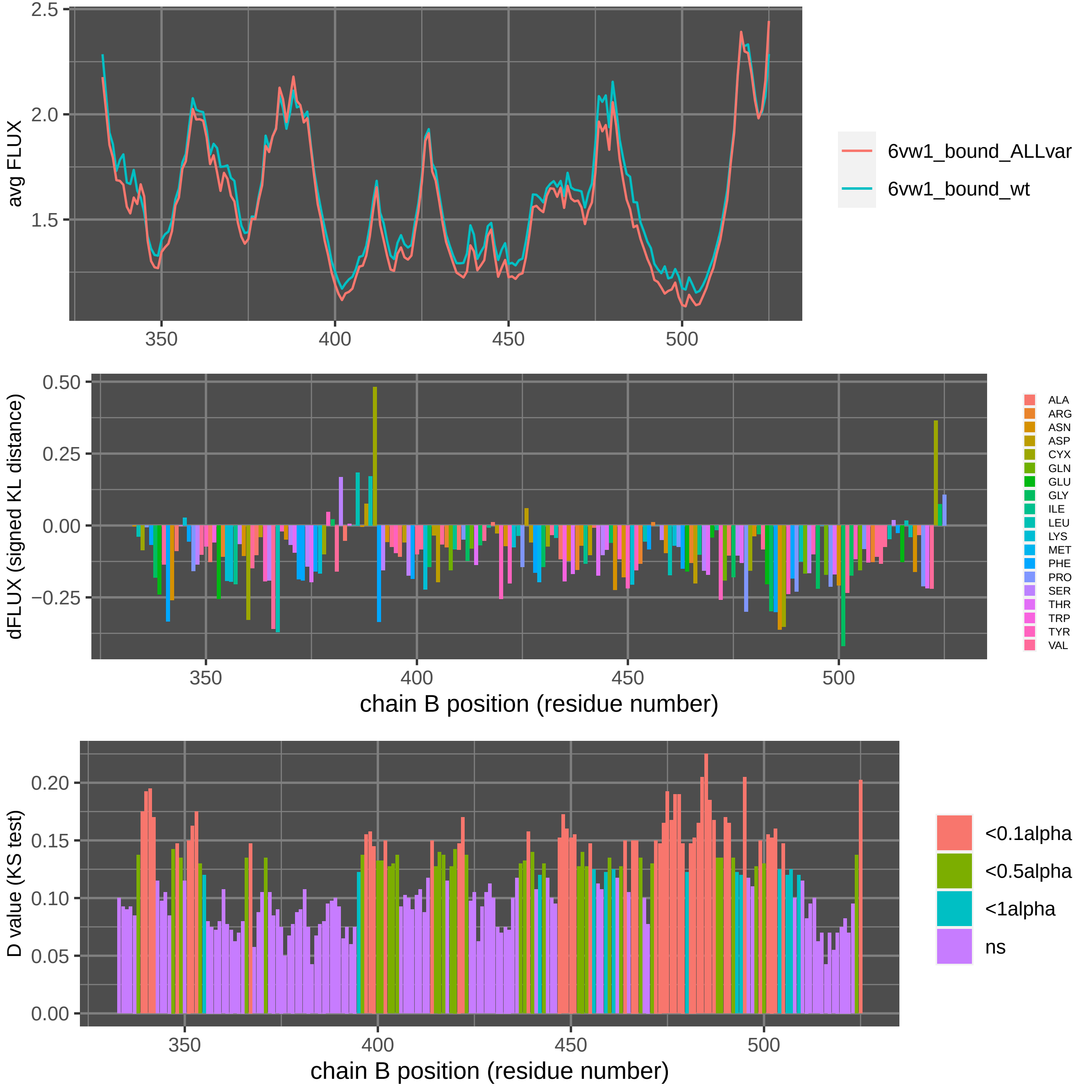

Supplement: Supplemental Data [file 471374_file09.zip › data_RajendranFerranBabbitt_2022/Figure 3D,3E [Sars-CoV-2 RBD WT_Mut(voc) + ACE2]/all_variants_ACE2bound/DROIDS_results_6vw1_bound_ALLvar_6vw1_bound_wt_flux_0.010_B/DROIDSplot_B.pdf]

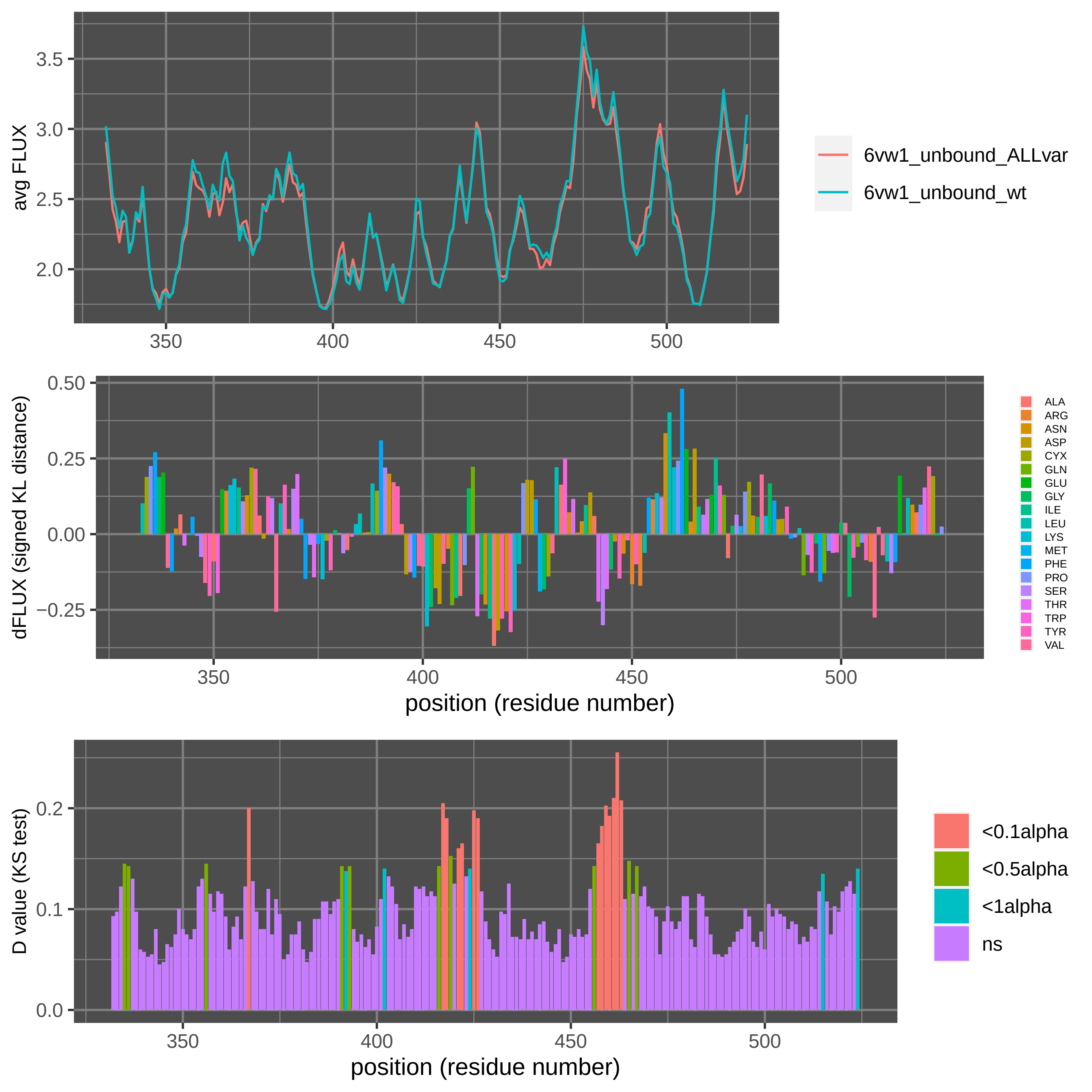

Supplement: Supplemental Data [file 471374_file09.zip › data_RajendranFerranBabbitt_2022/Figure 3D,3E [Sars-CoV-2 RBD WT_Mut(voc) + ACE2]/all_variants_ACE2unbound/DROIDS_results_6vw1_unbound_wt_6vw1_unbound_ALLvar_flux_0.010/DROIDSplot_all.pdf]

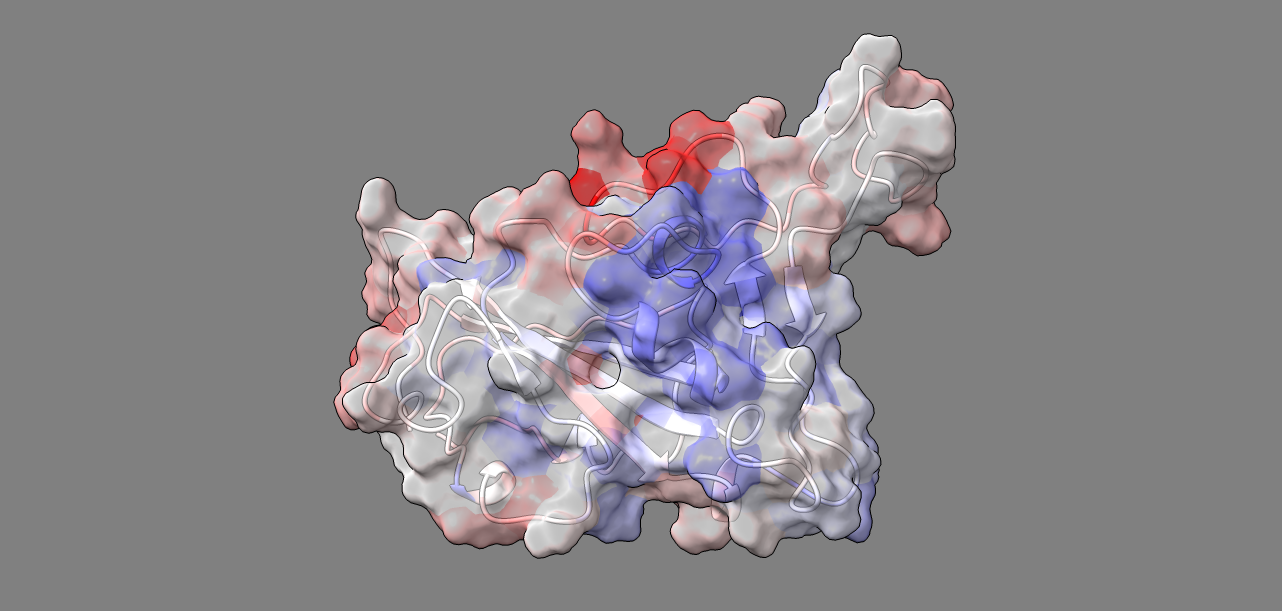

Supplement: Supplemental Data [file 471374_file09.zip › data_RajendranFerranBabbitt_2022/Figure 3D,3E [Sars-CoV-2 RBD WT_Mut(voc) + ACE2]/all_variants_ACE2unbound/DROIDS_results_6vw1_unbound_wt_6vw1_unbound_ALLvar_flux_0.010/image1.png]

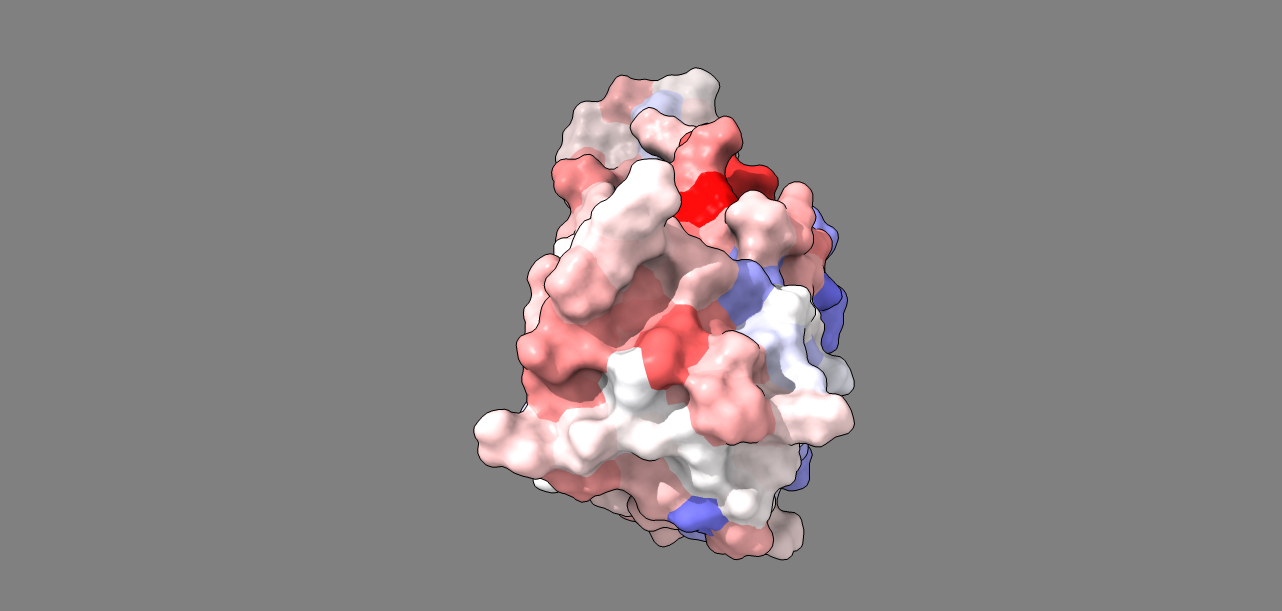

Supplement: Supplemental Data [file 471374_file09.zip › data_RajendranFerranBabbitt_2022/Figure 3D,3E [Sars-CoV-2 RBD WT_Mut(voc) + ACE2]/all_variants_ACE2unbound/DROIDS_results_6vw1_unbound_wt_6vw1_unbound_ALLvar_flux_0.010/image10.png]

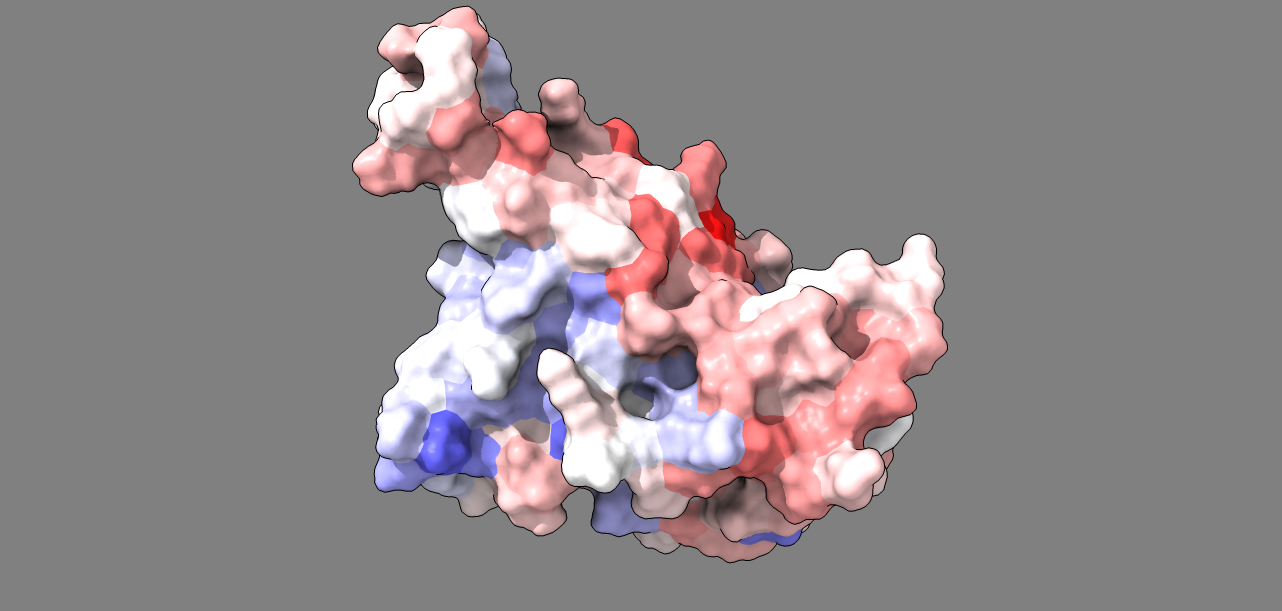

Supplement: Supplemental Data [file 471374_file09.zip › data_RajendranFerranBabbitt_2022/Figure 3D,3E [Sars-CoV-2 RBD WT_Mut(voc) + ACE2]/all_variants_ACE2unbound/DROIDS_results_6vw1_unbound_wt_6vw1_unbound_ALLvar_flux_0.010/image11.png]

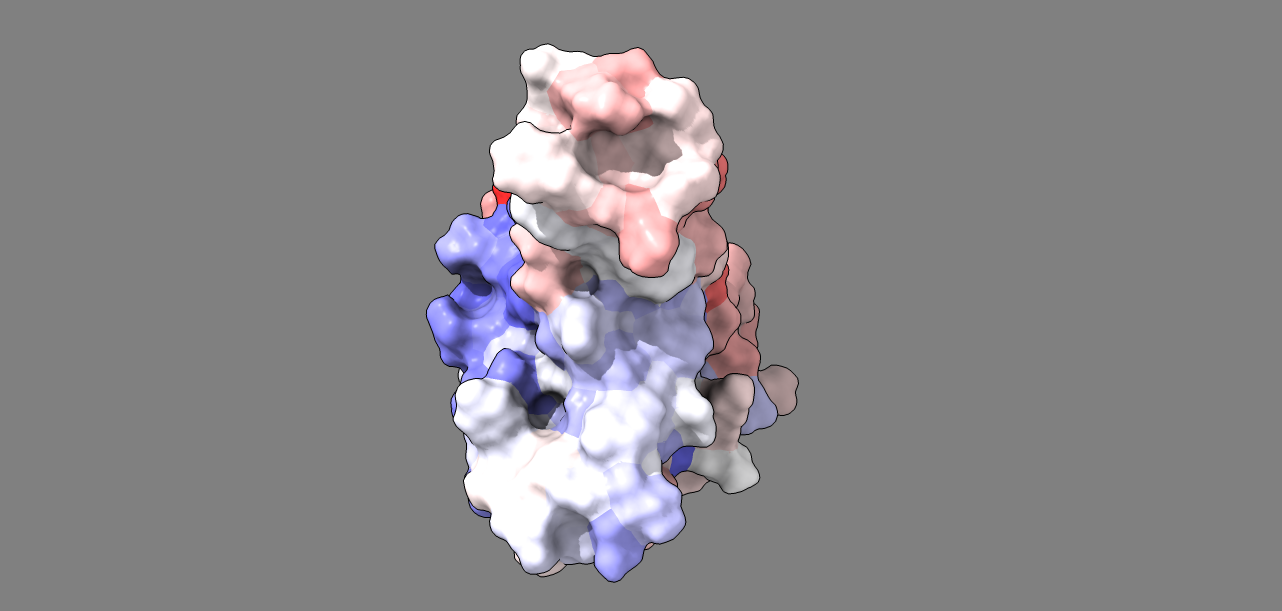

Supplement: Supplemental Data [file 471374_file09.zip › data_RajendranFerranBabbitt_2022/Figure 3D,3E [Sars-CoV-2 RBD WT_Mut(voc) + ACE2]/all_variants_ACE2unbound/DROIDS_results_6vw1_unbound_wt_6vw1_unbound_ALLvar_flux_0.010/image12.png]

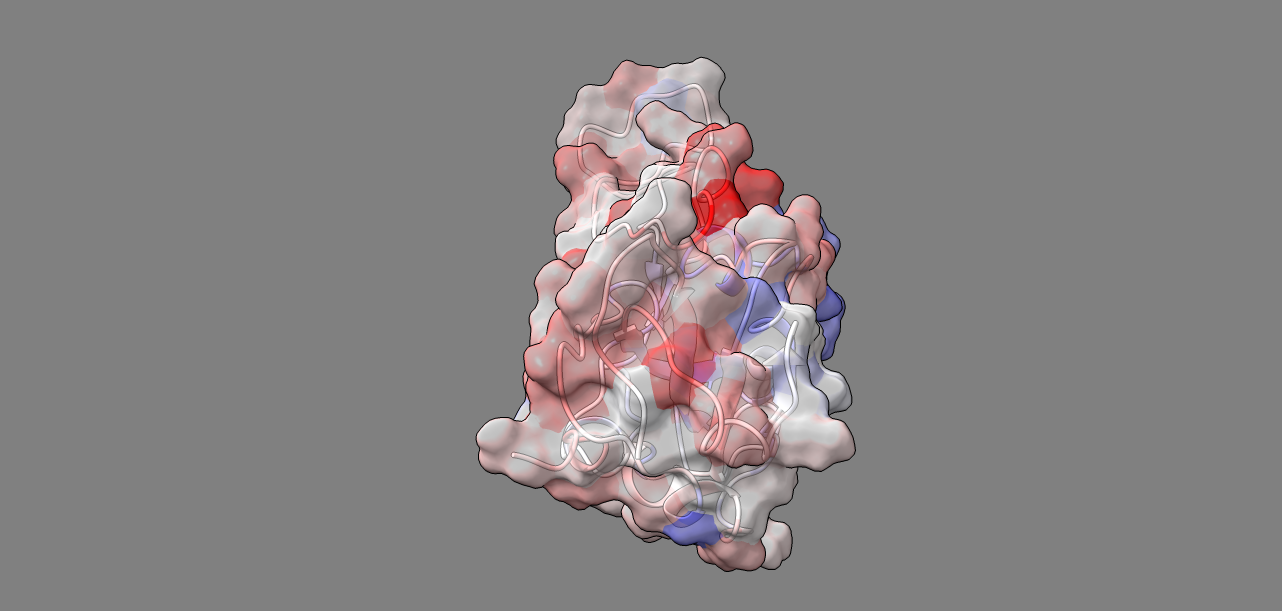

Supplement: Supplemental Data [file 471374_file09.zip › data_RajendranFerranBabbitt_2022/Figure 3D,3E [Sars-CoV-2 RBD WT_Mut(voc) + ACE2]/all_variants_ACE2unbound/DROIDS_results_6vw1_unbound_wt_6vw1_unbound_ALLvar_flux_0.010/image2.png]

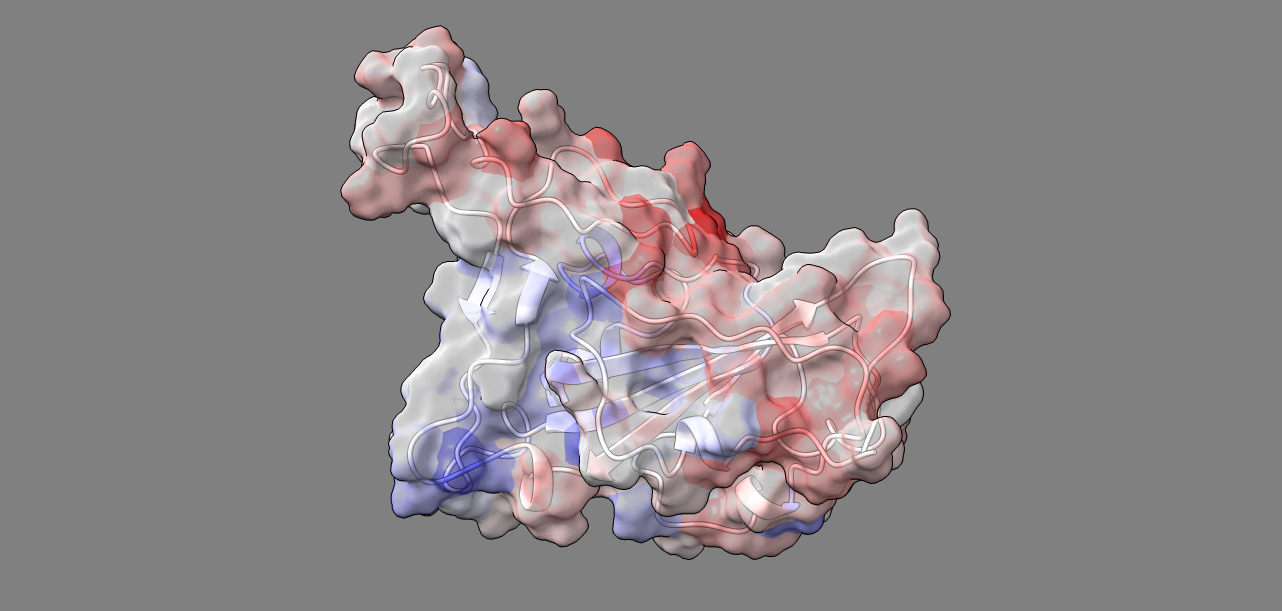

Supplement: Supplemental Data [file 471374_file09.zip › data_RajendranFerranBabbitt_2022/Figure 3D,3E [Sars-CoV-2 RBD WT_Mut(voc) + ACE2]/all_variants_ACE2unbound/DROIDS_results_6vw1_unbound_wt_6vw1_unbound_ALLvar_flux_0.010/image3.png]

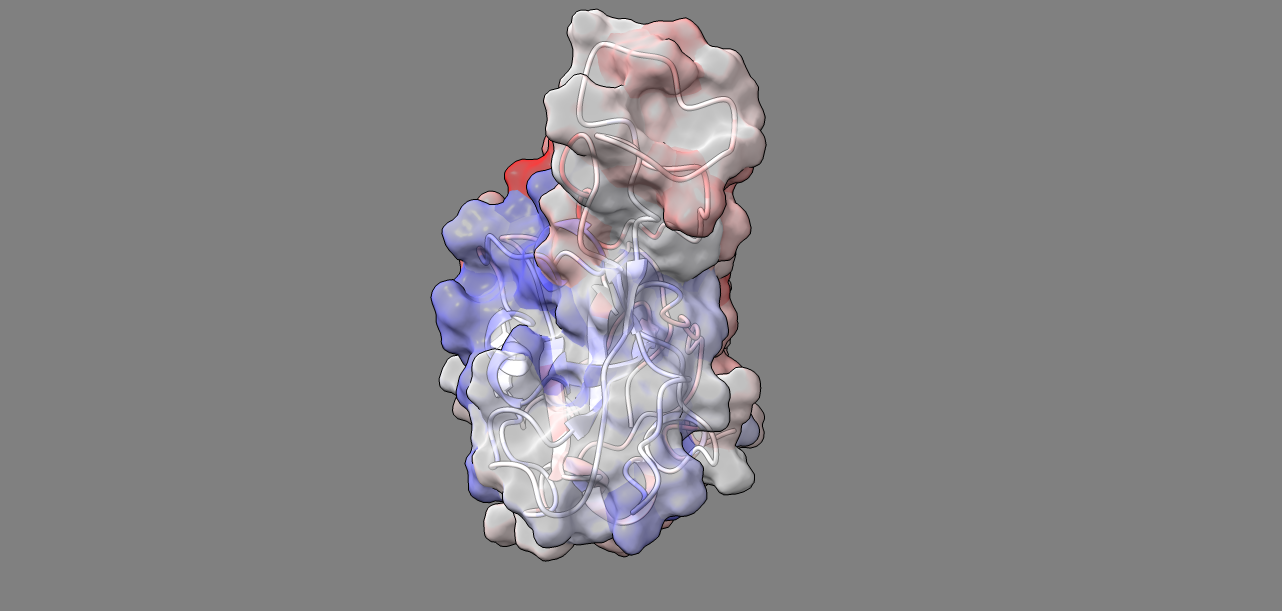

Supplement: Supplemental Data [file 471374_file09.zip › data_RajendranFerranBabbitt_2022/Figure 3D,3E [Sars-CoV-2 RBD WT_Mut(voc) + ACE2]/all_variants_ACE2unbound/DROIDS_results_6vw1_unbound_wt_6vw1_unbound_ALLvar_flux_0.010/image4.png]

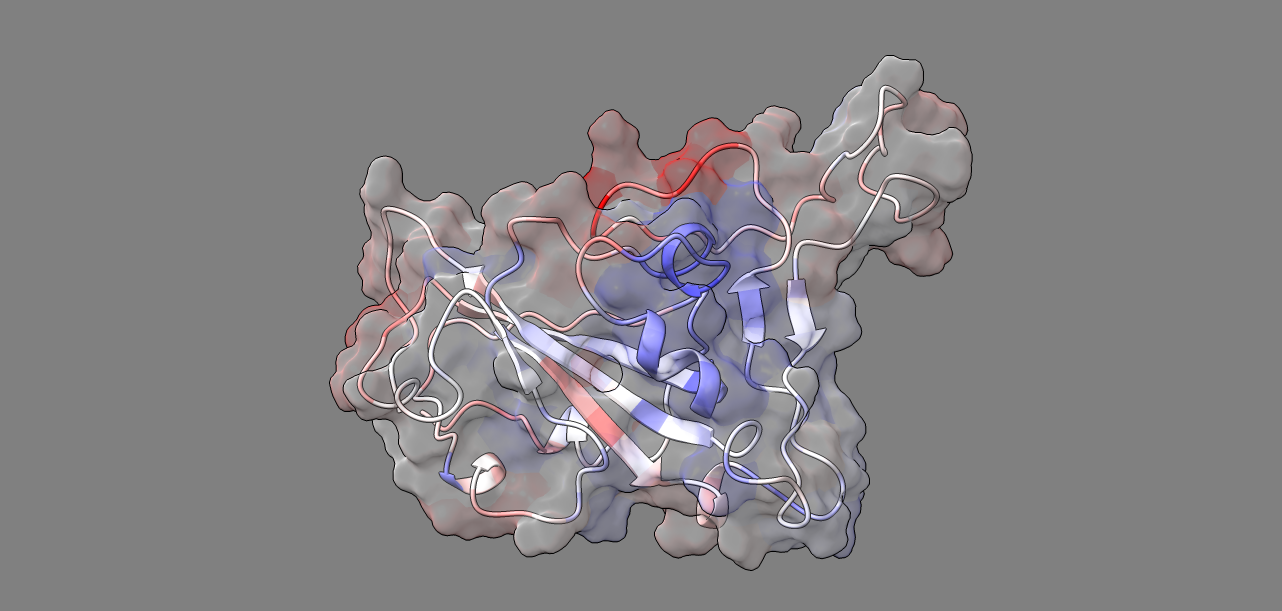

Supplement: Supplemental Data [file 471374_file09.zip › data_RajendranFerranBabbitt_2022/Figure 3D,3E [Sars-CoV-2 RBD WT_Mut(voc) + ACE2]/all_variants_ACE2unbound/DROIDS_results_6vw1_unbound_wt_6vw1_unbound_ALLvar_flux_0.010/image5.png]

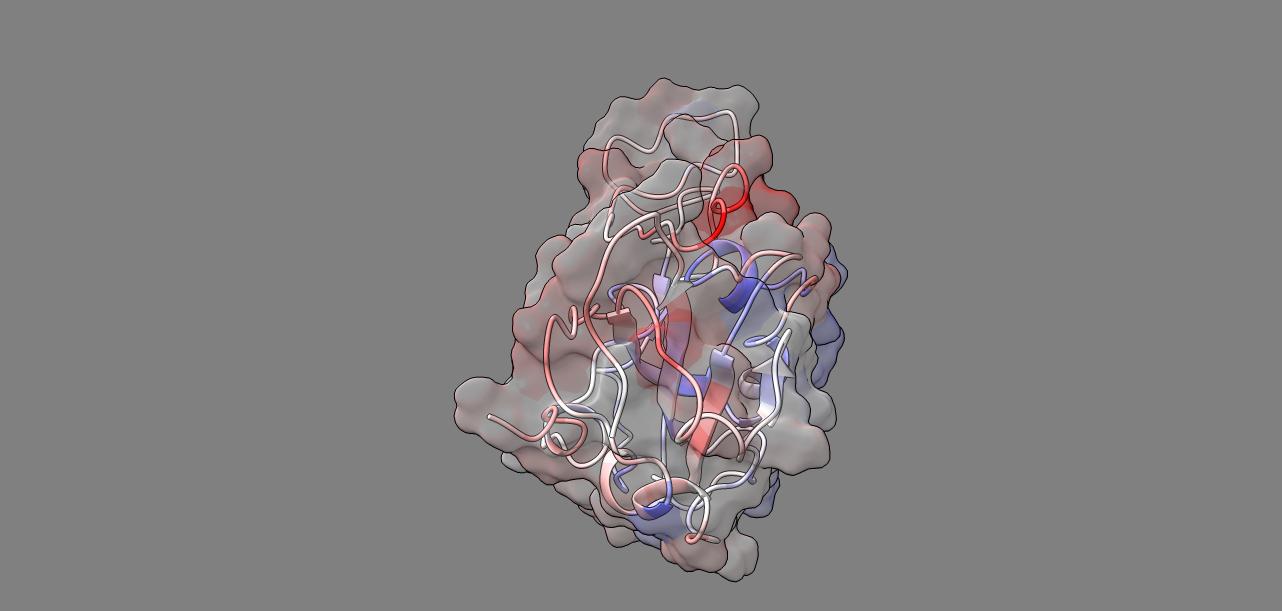

Supplement: Supplemental Data [file 471374_file09.zip › data_RajendranFerranBabbitt_2022/Figure 3D,3E [Sars-CoV-2 RBD WT_Mut(voc) + ACE2]/all_variants_ACE2unbound/DROIDS_results_6vw1_unbound_wt_6vw1_unbound_ALLvar_flux_0.010/image6.png]

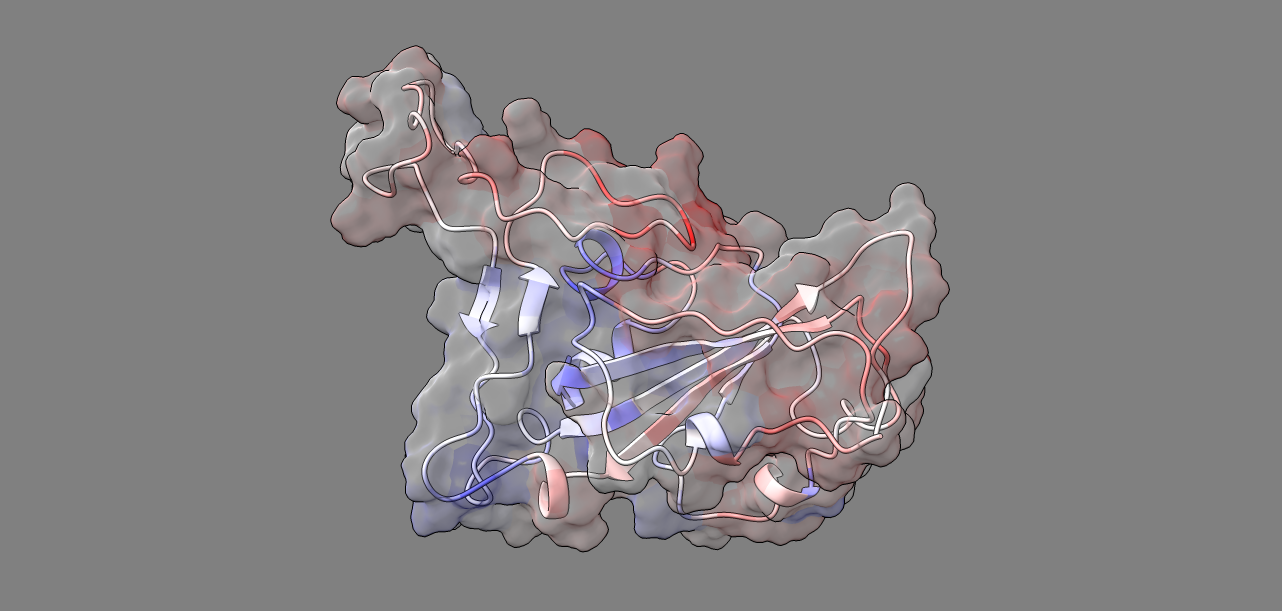

Supplement: Supplemental Data [file 471374_file09.zip › data_RajendranFerranBabbitt_2022/Figure 3D,3E [Sars-CoV-2 RBD WT_Mut(voc) + ACE2]/all_variants_ACE2unbound/DROIDS_results_6vw1_unbound_wt_6vw1_unbound_ALLvar_flux_0.010/image7.png]

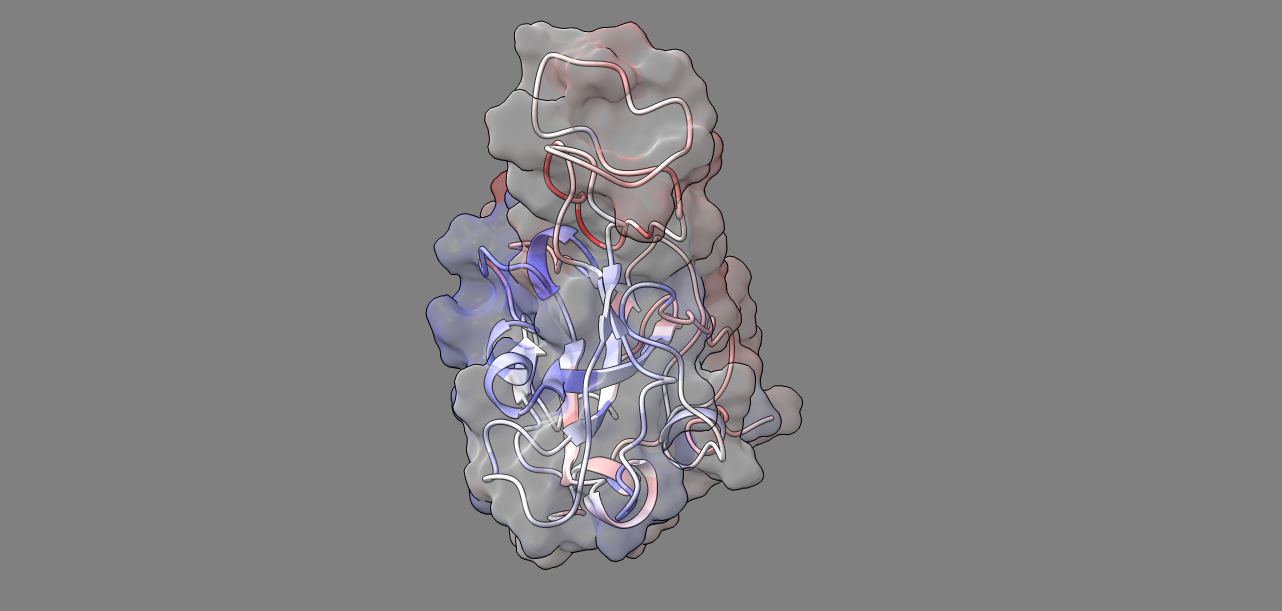

Supplement: Supplemental Data [file 471374_file09.zip › data_RajendranFerranBabbitt_2022/Figure 3D,3E [Sars-CoV-2 RBD WT_Mut(voc) + ACE2]/all_variants_ACE2unbound/DROIDS_results_6vw1_unbound_wt_6vw1_unbound_ALLvar_flux_0.010/image8.png]

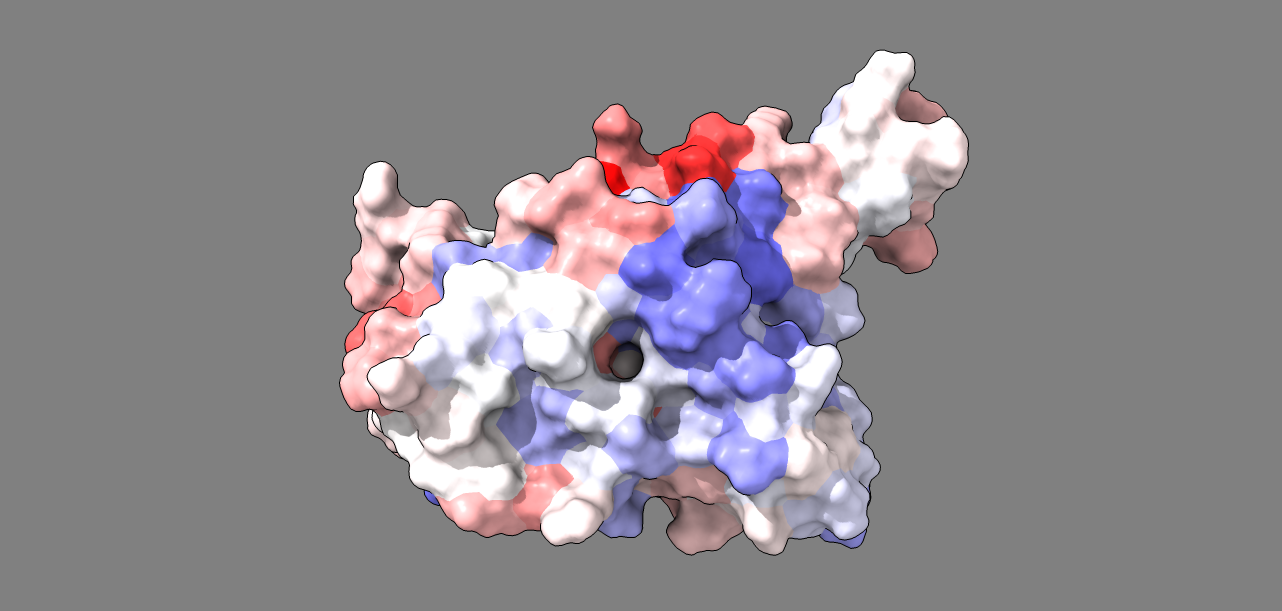

Supplement: Supplemental Data [file 471374_file09.zip › data_RajendranFerranBabbitt_2022/Figure 3D,3E [Sars-CoV-2 RBD WT_Mut(voc) + ACE2]/all_variants_ACE2unbound/DROIDS_results_6vw1_unbound_wt_6vw1_unbound_ALLvar_flux_0.010/image9.png]

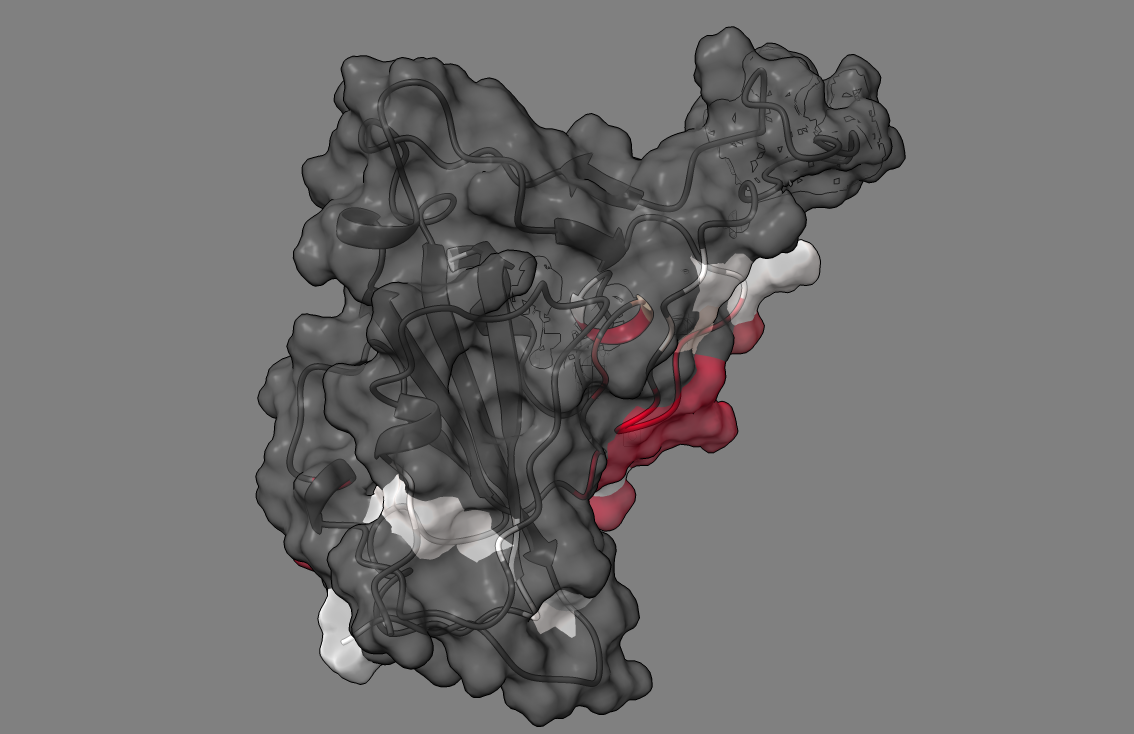

Supplement: Supplemental Data [file 471374_file09.zip › data_RajendranFerranBabbitt_2022/Figure 3D,3E [Sars-CoV-2 RBD WT_Mut(voc) + ACE2]/all_variants_ACE2unbound/DROIDS_results_6vw1_unbound_wt_6vw1_unbound_ALLvar_flux_0.010/p_values/image1.png]

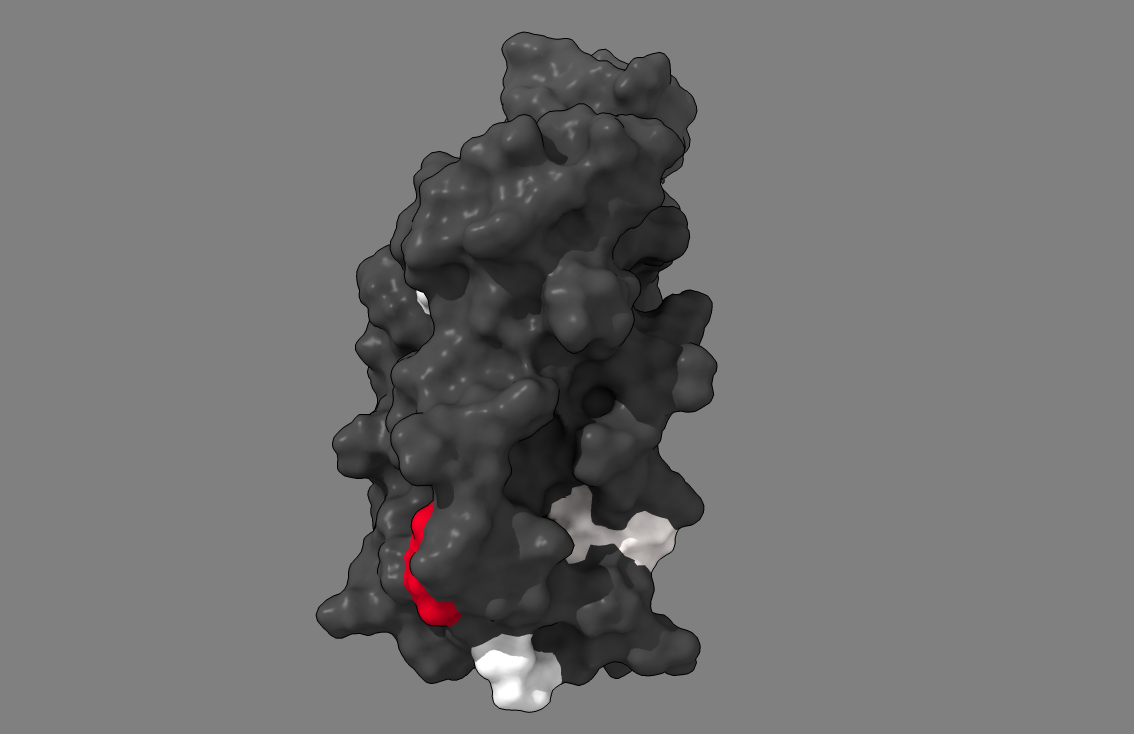

Supplement: Supplemental Data [file 471374_file09.zip › data_RajendranFerranBabbitt_2022/Figure 3D,3E [Sars-CoV-2 RBD WT_Mut(voc) + ACE2]/all_variants_ACE2unbound/DROIDS_results_6vw1_unbound_wt_6vw1_unbound_ALLvar_flux_0.010/p_values/image10.png]

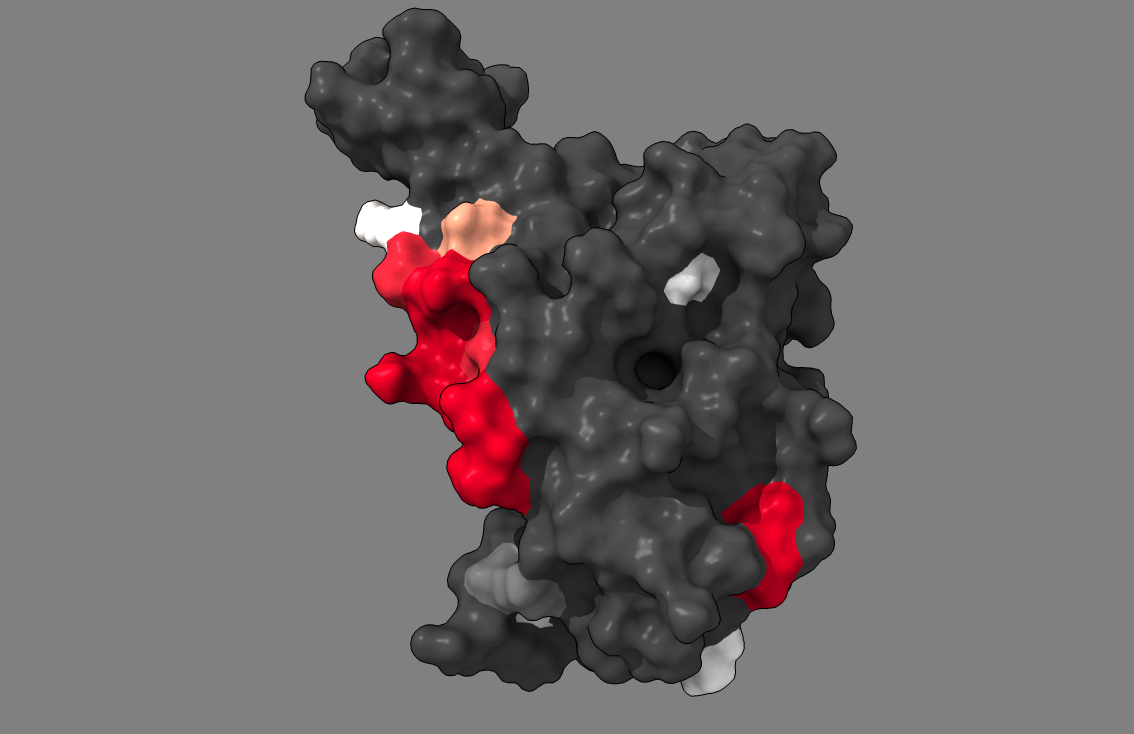

Supplement: Supplemental Data [file 471374_file09.zip › data_RajendranFerranBabbitt_2022/Figure 3D,3E [Sars-CoV-2 RBD WT_Mut(voc) + ACE2]/all_variants_ACE2unbound/DROIDS_results_6vw1_unbound_wt_6vw1_unbound_ALLvar_flux_0.010/p_values/image11.png]

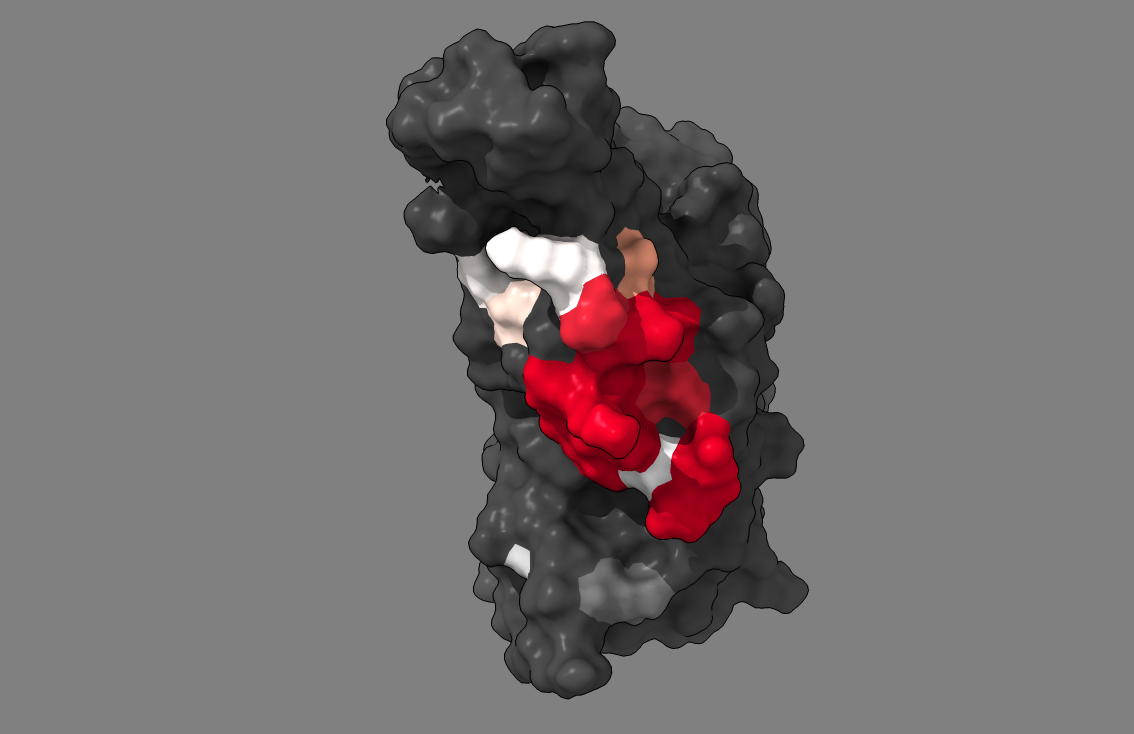

Supplement: Supplemental Data [file 471374_file09.zip › data_RajendranFerranBabbitt_2022/Figure 3D,3E [Sars-CoV-2 RBD WT_Mut(voc) + ACE2]/all_variants_ACE2unbound/DROIDS_results_6vw1_unbound_wt_6vw1_unbound_ALLvar_flux_0.010/p_values/image12.png]

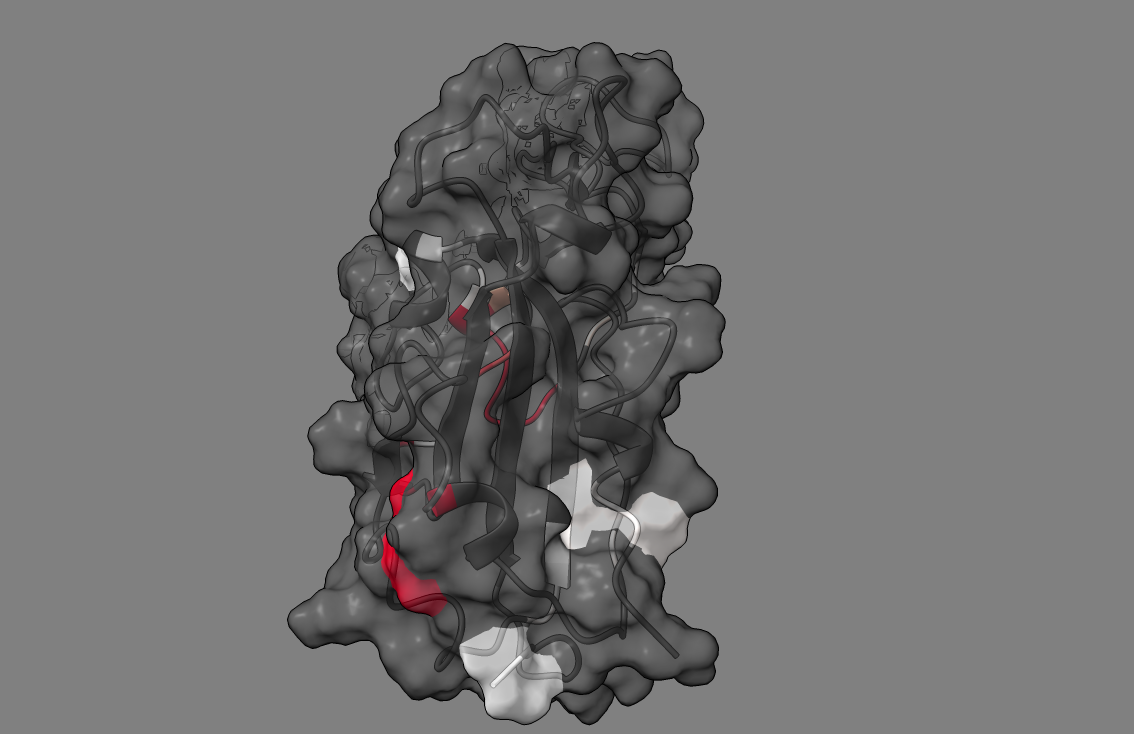

Supplement: Supplemental Data [file 471374_file09.zip › data_RajendranFerranBabbitt_2022/Figure 3D,3E [Sars-CoV-2 RBD WT_Mut(voc) + ACE2]/all_variants_ACE2unbound/DROIDS_results_6vw1_unbound_wt_6vw1_unbound_ALLvar_flux_0.010/p_values/image2.png]

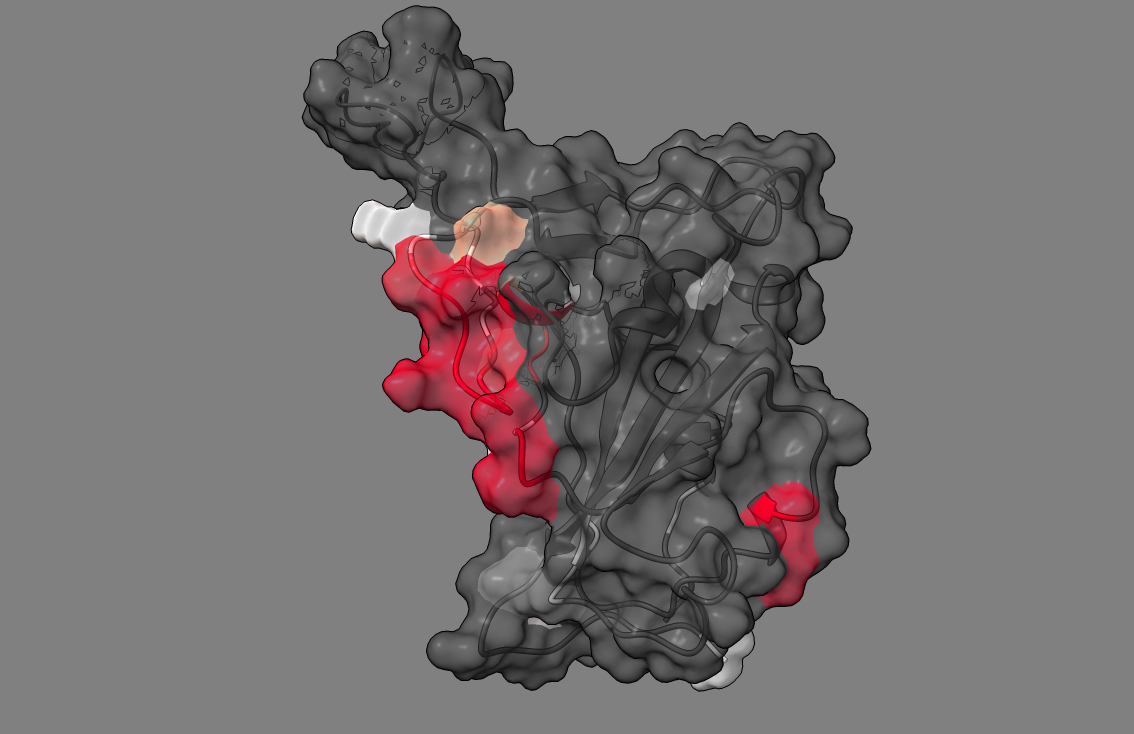

Supplement: Supplemental Data [file 471374_file09.zip › data_RajendranFerranBabbitt_2022/Figure 3D,3E [Sars-CoV-2 RBD WT_Mut(voc) + ACE2]/all_variants_ACE2unbound/DROIDS_results_6vw1_unbound_wt_6vw1_unbound_ALLvar_flux_0.010/p_values/image3.png]

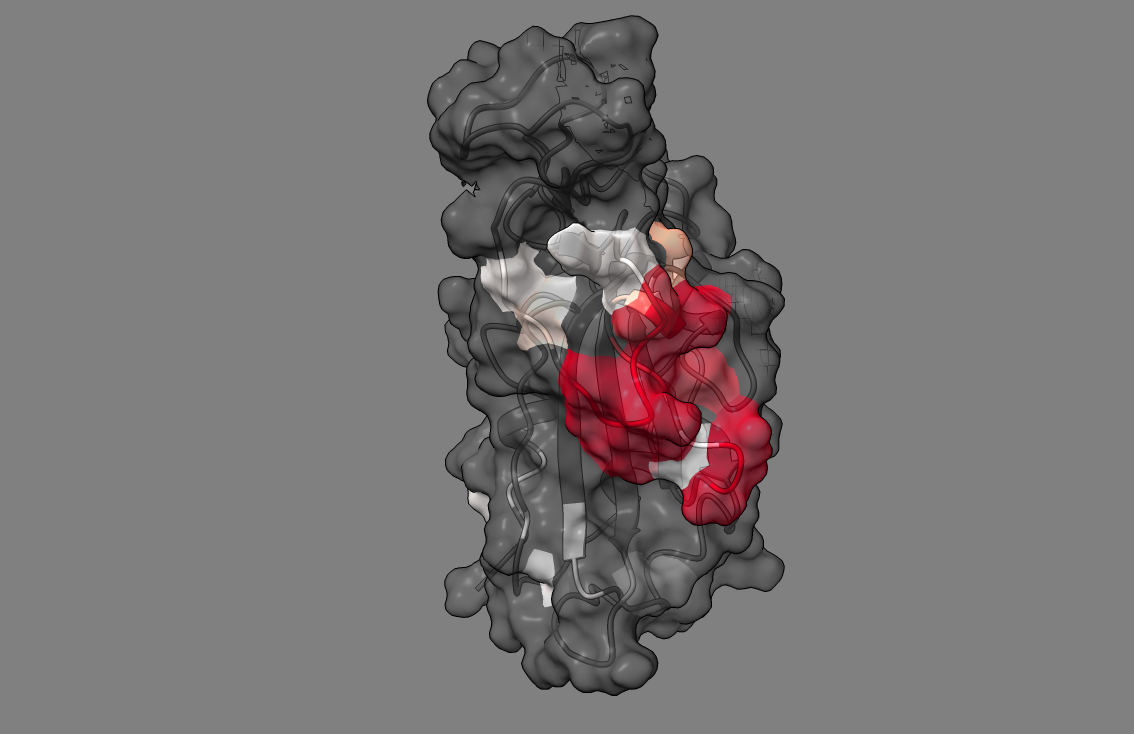

Supplement: Supplemental Data [file 471374_file09.zip › data_RajendranFerranBabbitt_2022/Figure 3D,3E [Sars-CoV-2 RBD WT_Mut(voc) + ACE2]/all_variants_ACE2unbound/DROIDS_results_6vw1_unbound_wt_6vw1_unbound_ALLvar_flux_0.010/p_values/image4.png]

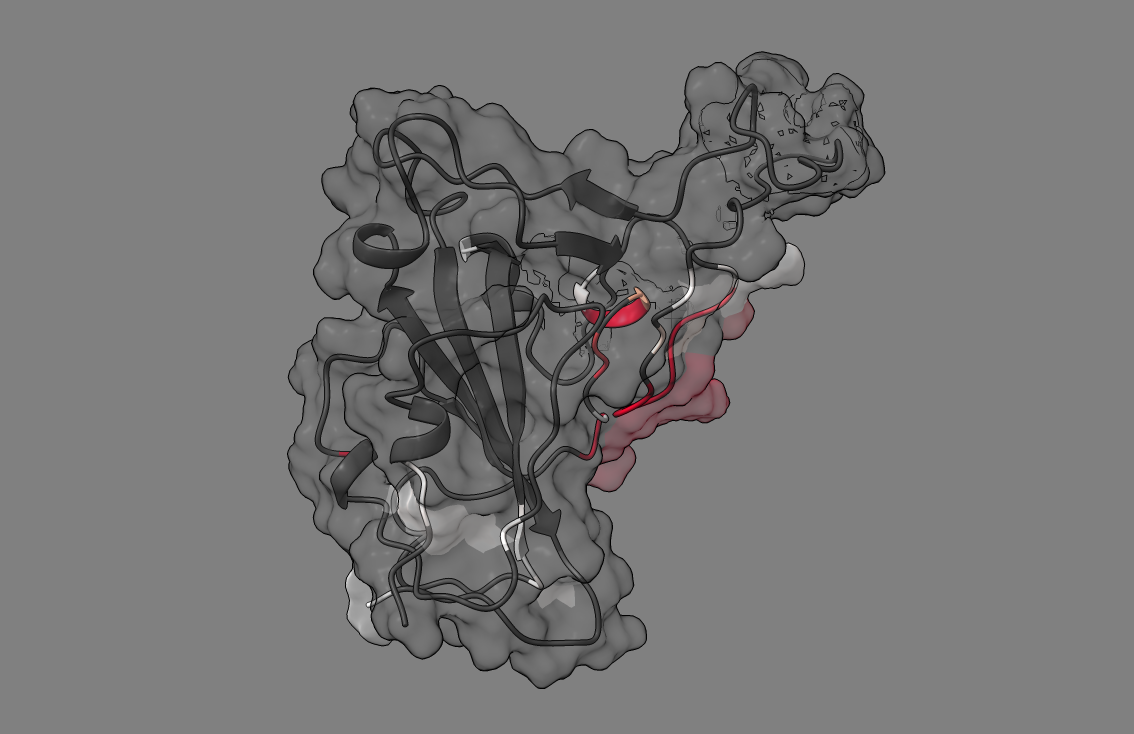

Supplement: Supplemental Data [file 471374_file09.zip › data_RajendranFerranBabbitt_2022/Figure 3D,3E [Sars-CoV-2 RBD WT_Mut(voc) + ACE2]/all_variants_ACE2unbound/DROIDS_results_6vw1_unbound_wt_6vw1_unbound_ALLvar_flux_0.010/p_values/image5.png]

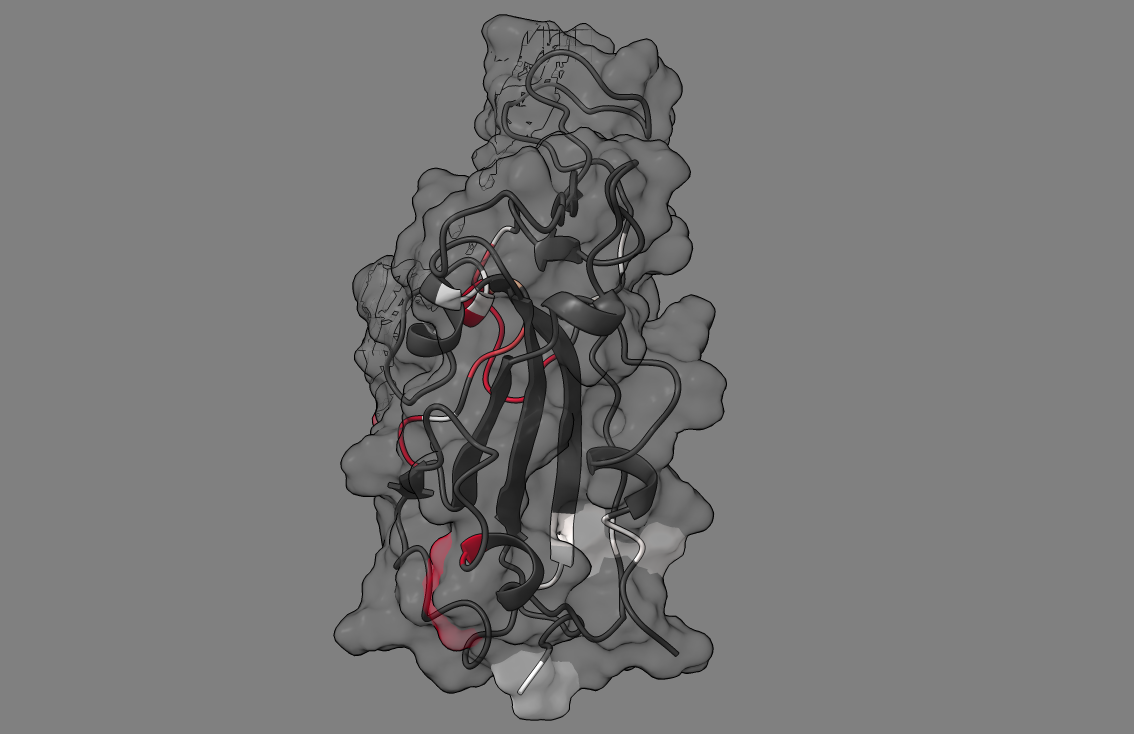

Supplement: Supplemental Data [file 471374_file09.zip › data_RajendranFerranBabbitt_2022/Figure 3D,3E [Sars-CoV-2 RBD WT_Mut(voc) + ACE2]/all_variants_ACE2unbound/DROIDS_results_6vw1_unbound_wt_6vw1_unbound_ALLvar_flux_0.010/p_values/image6.png]

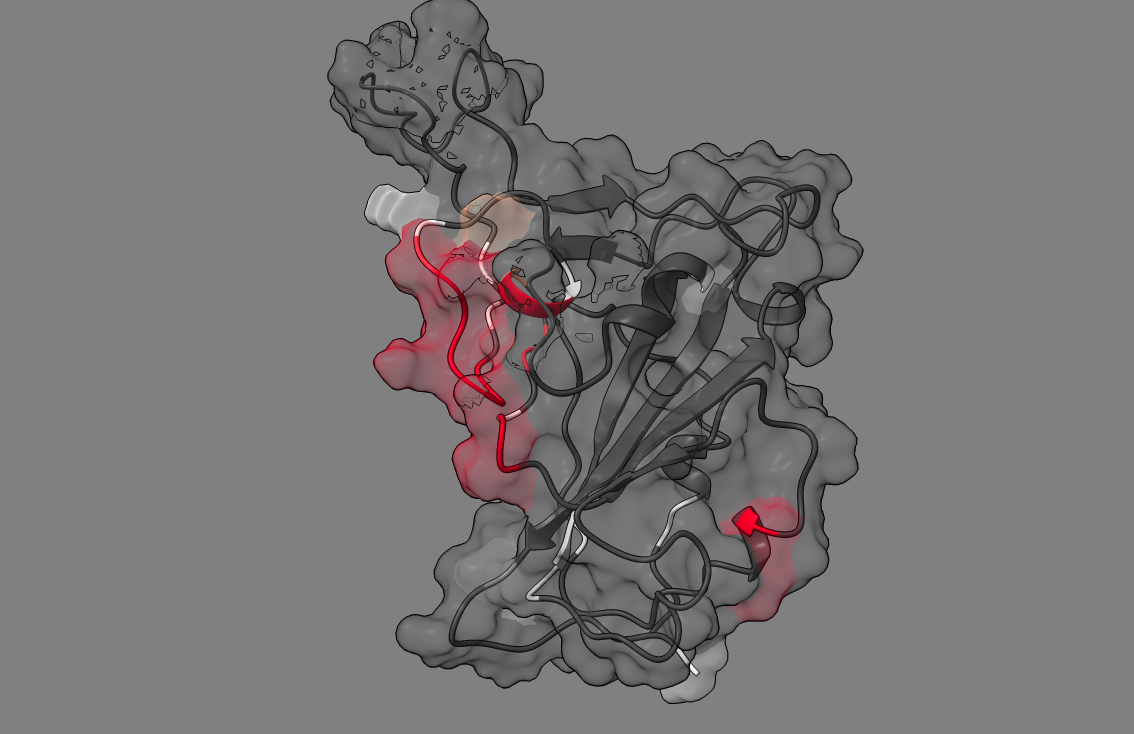

Supplement: Supplemental Data [file 471374_file09.zip › data_RajendranFerranBabbitt_2022/Figure 3D,3E [Sars-CoV-2 RBD WT_Mut(voc) + ACE2]/all_variants_ACE2unbound/DROIDS_results_6vw1_unbound_wt_6vw1_unbound_ALLvar_flux_0.010/p_values/image7.png]

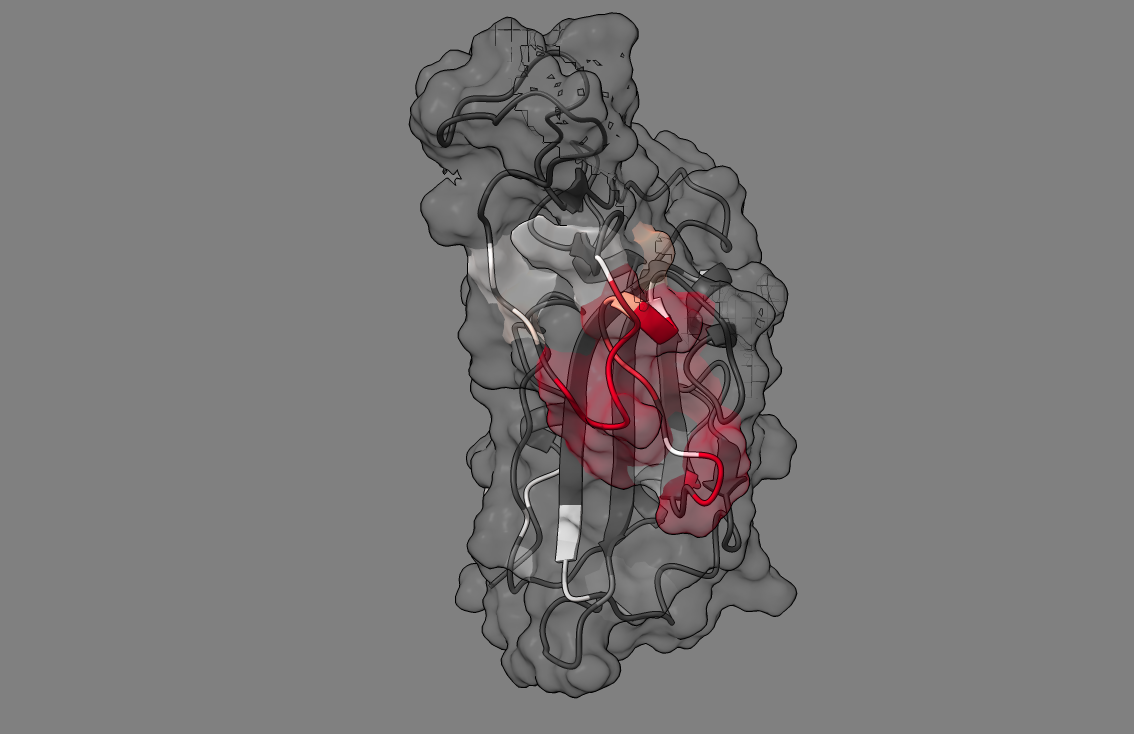

Supplement: Supplemental Data [file 471374_file09.zip › data_RajendranFerranBabbitt_2022/Figure 3D,3E [Sars-CoV-2 RBD WT_Mut(voc) + ACE2]/all_variants_ACE2unbound/DROIDS_results_6vw1_unbound_wt_6vw1_unbound_ALLvar_flux_0.010/p_values/image8.png]

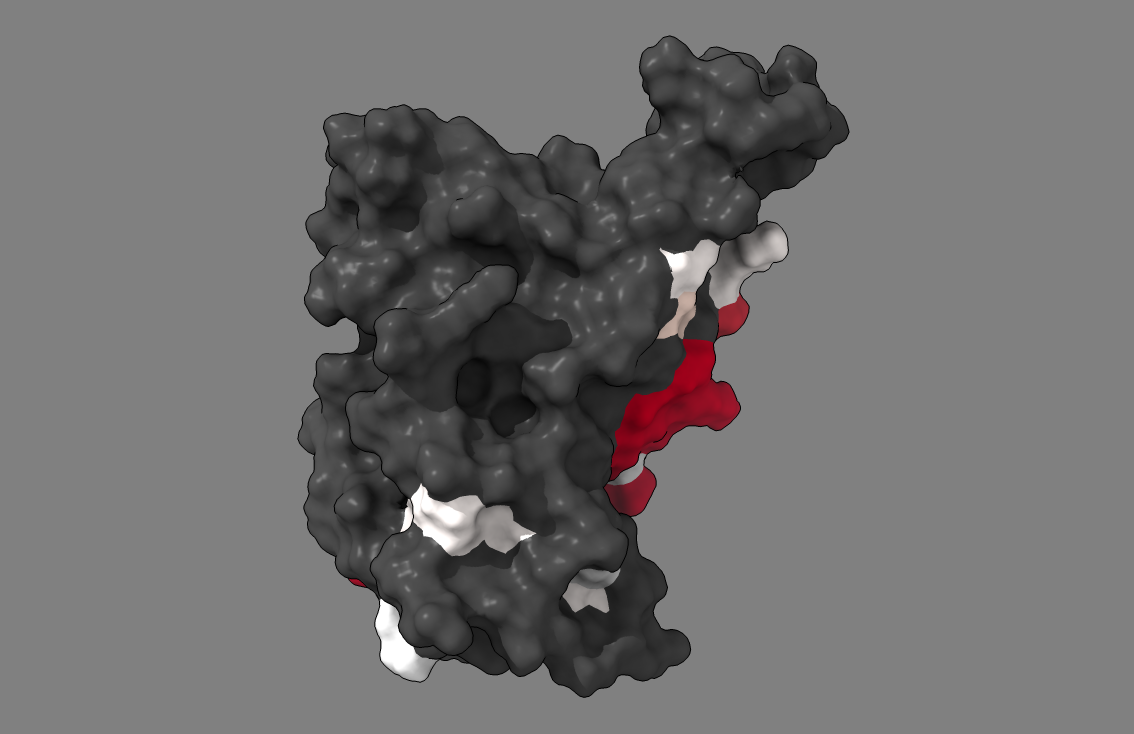

Supplement: Supplemental Data [file 471374_file09.zip › data_RajendranFerranBabbitt_2022/Figure 3D,3E [Sars-CoV-2 RBD WT_Mut(voc) + ACE2]/all_variants_ACE2unbound/DROIDS_results_6vw1_unbound_wt_6vw1_unbound_ALLvar_flux_0.010/p_values/image9.png]

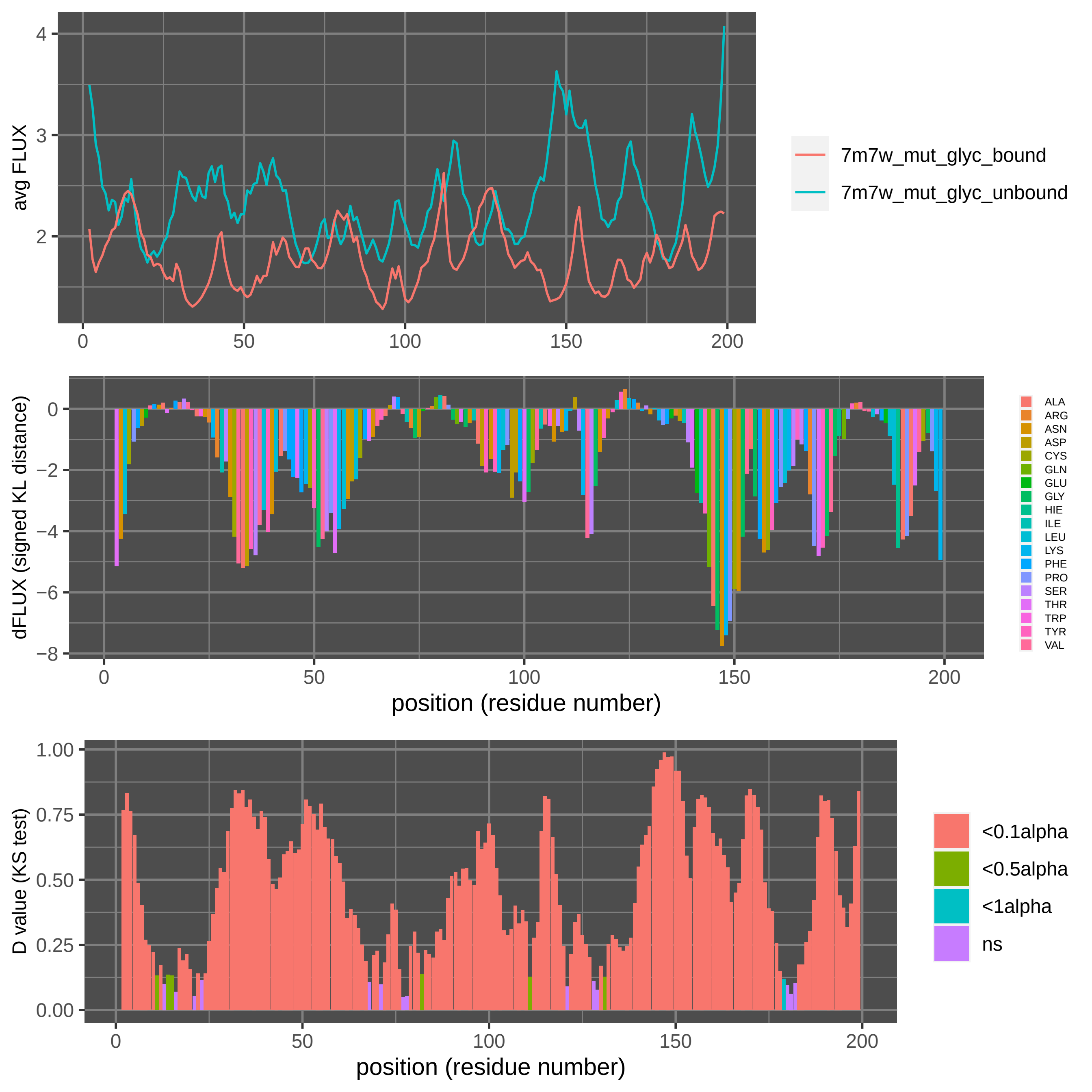

Supplement: Supplemental Data [file 471374_file09.zip › data_RajendranFerranBabbitt_2022/Figure 4,S1[omicron RBD + S2H97]/DROIDSplot_all.pdf]
